# Generation of T cells with reduced off-target cross-reactivities by engineering co-signalling receptors

**DOI:** 10.1101/2024.10.25.620274

**Authors:** Jose Cabezas-Caballero, Anna Huhn, Mikhail A. Kutuzov, Violaine Andre, Alina Shomuradova, P. Anton van der Merwe, Omer Dushek

## Abstract

Adoptive T cell therapy using T cells engineered with novel T cell receptors (TCRs) targeting tumor-specific peptides is a promising immunotherapy. However, these TCR-T cells can cross-react with off-target peptides, leading to severe autoimmune toxicities. Current efforts focus on identifying TCRs with reduced cross-reactivity. Here, we show that T cell cross-reactivity can be controlled by the co-signalling molecules CD5, CD8, and CD4, without modifying the TCR. We find the largest reduction in cytotoxic T cell cross-reactivity by knocking out CD8 and expressing CD4. Cytotoxic T cells engineered with a CD8-to-CD4 co-receptor switch show reduced cross-reactivity to random and positional scanning peptide libraries, as well as to self-peptides, while maintaining their on-target potency. Therefore, co-receptor switching generates super selective T cells that reduce the risk of lethal off-target cross-reactivity, and offers a universal method to enhance the safety of T cell immunotherapies for any TCR.

**Graphical abstract:** 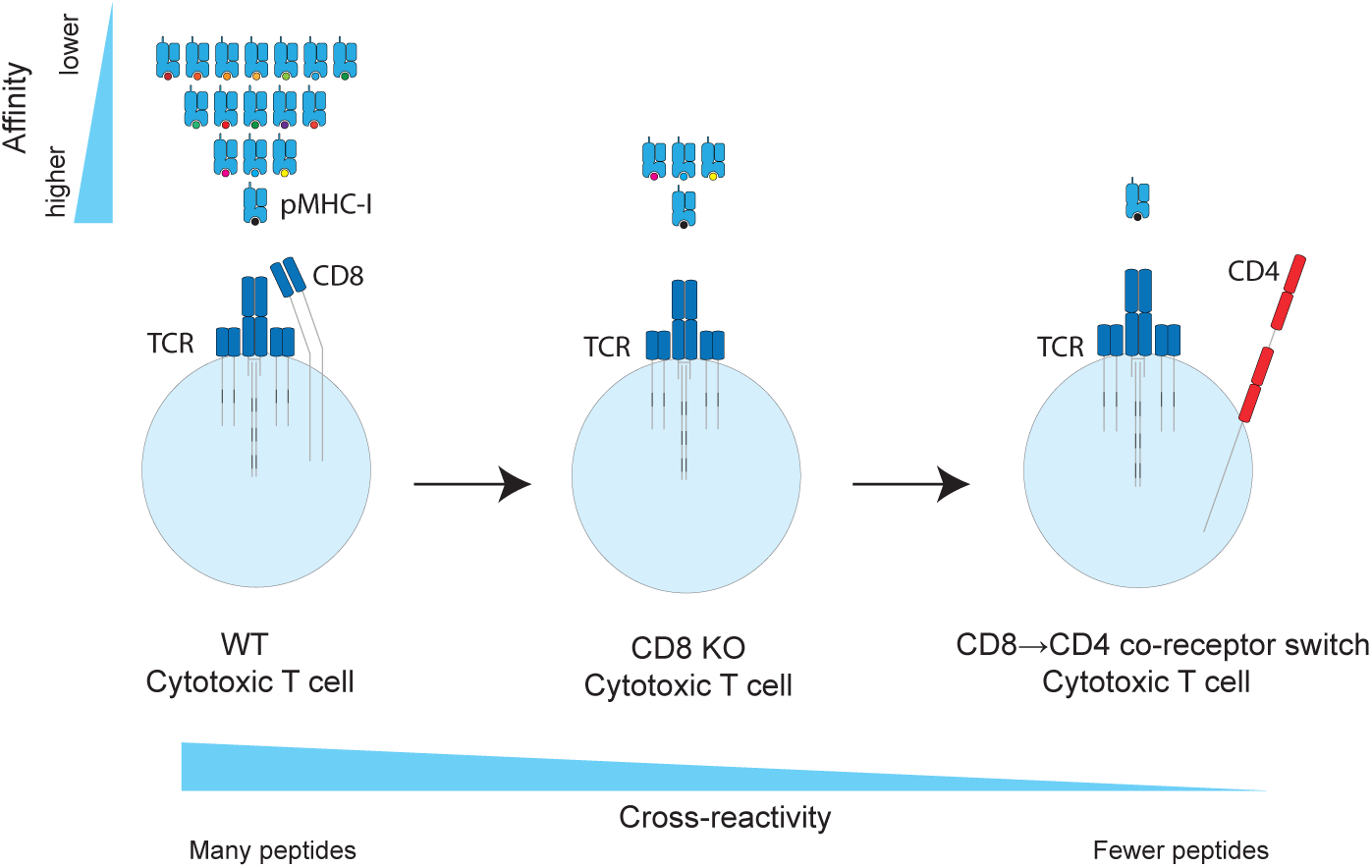

**One sentence summary:** Switching the CD8 for the CD4 co-receptor in cytotoxic T cells reduces the functional cross-reactivity of T cells without modifying the TCR.

## Introduction

A promising immunotherapy approach is the adoptive transfer of T cells engineered with novel T cell receptors (TCR-T) recognising tumour peptide antigens displayed on major histocompatibility complexes (pMHCs) (1). This therapeutic strategy enables targeting nearly all tumour antigens, including tumour specific developmental antigens and neo-antigens (2). However, the engineered T cells can cross-react with off-target peptides in healthy tissues and cause fatal autoimmune toxicities (3–5). This cross-reactivity has hampered efforts to produce highly potent TCR-T cell therapies (6, 7).

Identifying the potential off-target cross-reactivities of TCRs before first-in-human clinical trials is challenging due to the lack of animal models or cell lines that represent the entire human proteome and HLA allele diversity. Indeed, the clinical a3a TCR targeting the cancer-testis antigen MAGE-A3 passed safety screens but ultimately cross-reacted with a lower affinity off-target peptide from the cardiac protein Titin, causing the death of two patients (4, 5). As a result, efforts are underway to establish pipelines to identify effective yet safe TCRs (8–15). Typically, these methods screen TCRs with different complementary determining regions (CDRs) for their ability to recognise the on-target tumour but not off-target self pMHCs (16). In addition to screening methods, it has also been proposed that modifying the CDR loops to reduce their flexibility or introduce catch bonds may generally increase TCR specificity (17–19). However, these strategies that rely on mutating the TCR sequence to reduce cross-reactivity require prior knowledge of the self antigen that causes lethal cross-reactivity and modifying the TCR sequence to reduce cross-reactivity to one antigen may result in new cross-reactivities to other self antigens. Collectively, this makes it challenging and costly to screen and optimise each new candidate therapeutic TCR.

Instead of modifying the TCR CDR loops to reduce binding cross-reactivity, we hypothesised that functional cross-reactivity can be reduced by manipulating T cell signalling without modifying the TCR. In this way, even though the TCR can bind a large number peptides, T cells would only become activated in response to the few peptides that bind with high affinity. Put differently, we suggest that enhancing the ability of T cells to discriminate antigens based on their affinity would reduce their functional cross-reactivity. Given that co-signalling receptors on the T cell surface are known to impact TCR signalling (20), we reasoned that they impact T cell cross-reactivity.

Here, we established a platform to quantify the impact of co-signalling receptors on T cell ligand discrimination. While a knockout of the surface molecule CD5 decreased antigen discrimination, we found that a knockout of CD8 or expression of CD4 increased it. The largest effect was observed by combining CD8 knockout and CD4 expression (’co-receptor switch’). We demonstrate that a CD8→CD4 co-receptor switch dramatically reduced T cell cross-reactivity to peptide libraries and self peptides. Overall, co-receptor switching is a broadly applicable strategy to produce super selective T cells that minimise the risk of lethal cross-reactivities without compromising on-target potency, and can be applied to any TCR.

## Results

### T cell co-signalling receptors differentially modulate ligand sensitivity and discrimination

We established a platform to quantify the contribution of different T cell co-signalling receptors to ligand discrimination. We selected the NY-ESO-1 specific c259 TCR contained in the investigational TCR-T therapy lete-cel as a model system (21). First, we measured the binding affinity of the c259 TCR to a panel of 7 NY-ESO-1 peptide variants on HLA-A*02:01 by Surface Plasmon Resonance (SPR) (22) (Fig. S1, Table S1). Second, we used CRISPR/Cas9 to knock-out out five co-signalling receptors in primary human T cells expressing the c259 TCR that were previously suggested to impact ligand discrimination: CD8 (23), CD5 (24), CD43 (25), CD2 and LFA-1 (22, 26) (Fig. 1A). Third, we co-cultured these T cells with antigen-presenting-cells (APCs) pulsed with a titration of each of the 7 peptides with different affinities to the TCR and assessed their ability to induce multiple measures of T cell activation (target cell killing, IFN*γ* secretion, and 4-1BB upregulation). Finally, we quantified pMHC potency as the concentration of peptide required to elicit 15% activation (P15) from WT or KO T cells. By plotting the fold-change in potency (ΔP15) over affinity we could determine whether the co-sigalling molecule was selectively decreasing activation to lower-affinity ligands (Fig. 1B).

**Figure 1:**
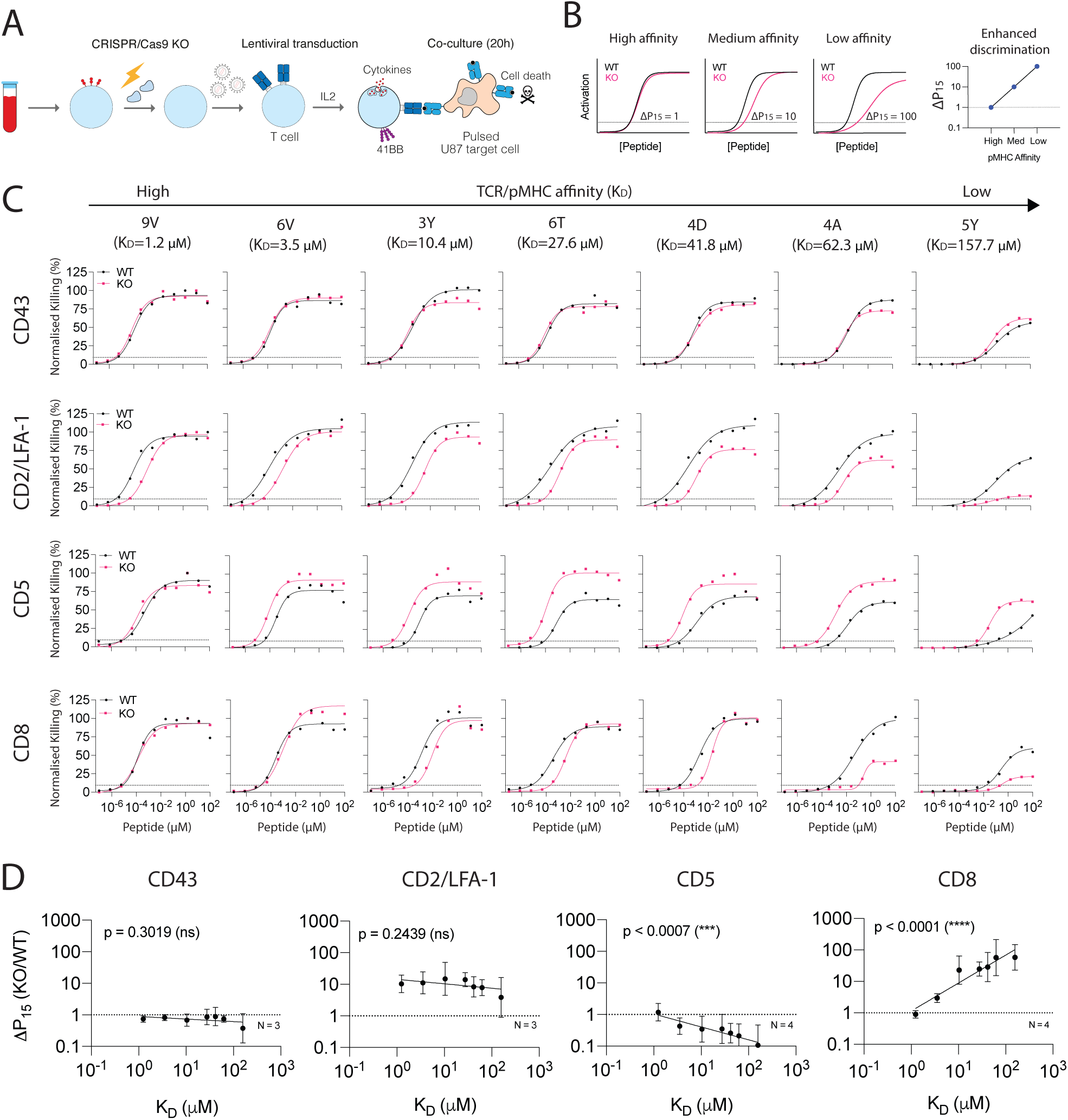
Measuring the impact of different T cell co-signalling receptors on ligand sensitivity and discrimination. **(A)** Experimental workflow to produce gene knockout primary human TCR-T cells. **(B)** Schematic of analysis method to determine the impact of gene knockout on ligand discrimination: changes in ligand potency between WT and KO TCR-T cells are plotted for different ligand affinities. Ligand potency (P15) is the ligand concentration required to activate 15% of maximum response. **(C)** U87 cells were titrated with each of the 7 NY-ESO-1 peptides to stimulate WT or KO c259 TCR-T cells. Killing of the target U87 cells was measured after 20 hours. Dashed line indicates potency (P15). **(D)** Fold change in potency (P15) between KO and WT T cells from (C) plotted over the TCR/pMHC affinity (K_D_). Dashed line indicates fold change of 1. Data in (C) are representative of at least N=3 independent experiments with different blood donors. Data in (D) is shown as means ± SDs. Significance of non-zero slope was assessed by an F-test.

We achieved high knockout efficiency of each co-signalling receptor (Fig. S2A) enabling assessment of their impact on ligand discrimination (Fig. 1C-D, S2-4). The knock-out of CD43 had no impact on activation whereas the knock-out of CD2 or LFA-1 individually or in combination reduced activation for all pMHC affinities to a similar extent and therefore, these molecules do not impact ligand discrimination. In contrast, a knock-out of CD5 selectively improved activation against lower affinity ligands and therefore, CD5 KO reduced ligand discrimination. The knockout of CD8 selectively reduced activation to lower affinity peptides without impacting the higher-affinity NY-ESO-1 target antigen and therefore, CD8 KO increases ligand discrimination. Since the c259 TCR is affinity-matured (27), we confirmed that CD8 KO also increased the discrimination of the parental wild-type 1G4 TCR (28) (Fig. S5). Taken together, co-signalling molecules can control TCR ligand discrimination and a CD8 KO in particular can selectively reduce activation to lower affinity ligands without impacting potency to the higher-affinity on-target antigen.

### CD8 knock-out abolishes therapeutic a3a TCR cross-reactivity to Titin

T cells engineered with the MAGE-A3 specific a3a TCR caused lethal cardiac toxicities in a clinical trial due to cross-reactivity to a lower affinity peptide from the muscle protein Titin (4, 5). Since we have demonstrated that the CD8 co-receptor can decrease T cell ligand discrimination, we decided to investigate whether the cross-reactivity to Titin was CD8 dependent.

Given that TCR-T therapies rely on expressing the therapeutic MHC-I restricted TCR in both CD8+ cytotoxic and CD4+ helper T cells, we first examined their individual abilities to react to each antigen. Whilst both cytotoxic and helper T cells responded to the on-target MAGE-A3 antigen, only cytotoxic T cells responded to the off-target Titin antigen confirming that cytotoxic T cells are the likely source of autoimmune toxicity (Fig. 2A-B). By knocking out CD8 in cytotoxic cells we abolish activation against Titin without impacting responses to the higher-affinity on-target antigen (Fig. 2C-D). The CD8 KO also abolished the activation of T cells against Nalm6 cells that endogenously express Titin (5) (Fig. 2E).

**Figure 2:**
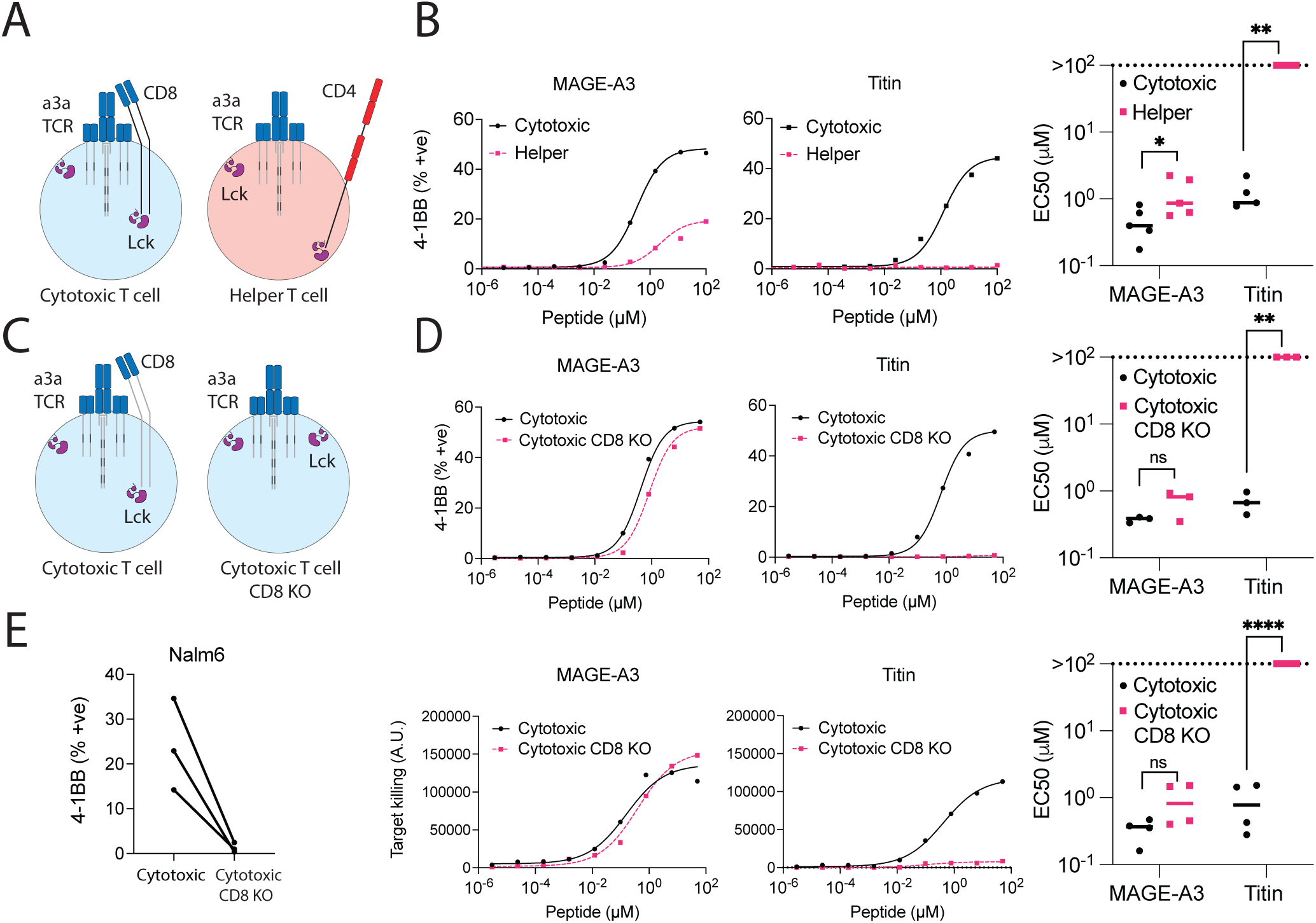
CD8 co-receptor KO abolishes MAGE-A3 TCR cross-reactivity to the self-antigen Titin. **(A)** Schematic of helper and cytotoxic T cells transduced with the MAGE-A3 specific a3a TCR. Lck can exist in a free state or a co-receptor bound state. **(B)** HLA-A1+ T2 cells were titrated with MAGE-A3 or Titin peptides to stimulate cytotoxic or helper a3a TCR-T cells for 20 hours. Representative dose-responses (Left) and mean sensitivity as EC50 (Right). **(C)** Schematic of WT or CD8 KO cytotoxic a3a TCR-T cells. Lck can exist in a free state or a co-receptor bound state. **(D)** HLA-A1+ T2 cells were titrated with MAGE-A3 or Titin peptides to stimulate WT or CD8 KO cytotoxic a3a TCR-T cells for 20 hours. Representative dose-responses (Left) and mean sensitivity as EC50 (Right). Data measuring 4-1BB surface activation marker (top) and target cell killing (bottom) are shown. **(E)** Nalm6 cells endogenously expressing the Titin protein were co-cultured with WT or CD8 KO cytotoxic a3a TCR-T cells for 20 hours. 4-1BB was stained by flow cytometry. Each data point in (E) represents an independent experiment with different blood donors. Each EC50 data point in (B) and (D) represents an independent experiment with different blood donors. P values were determined by paired t-test; ns not significant, *p*<*0.05, **p*<*0.01, ****p*<*0.0001

### Helper T cells display enhanced discrimination against pMHC-I antigens due to their incompatible CD4 co-receptor

The observation that CD4+ helper T cells only responded to the higher-affinity MAGE-A3 antigen whereas CD8+ cytotoxic T cells also responded to the lower-affinity Titin antigen (Fig. 2A-B) suggested that helper T cells may have a different capacity to discriminate ligands.

We compared ligand discrimination in helper vs cytotoxic T cells using the NY-ESO-1 c259 TCR platform (Fig. 3A). Consistent with the a3a TCR, we found that cytotoxic T cells activated more strongly against low affinity pMHCs than helper T cells (Fig. 3B-C). Interestingly, helper T cells displayed even higher discrimination than CD8 KO cytotoxic T cells (Fig. 3C).

**Figure 3:**
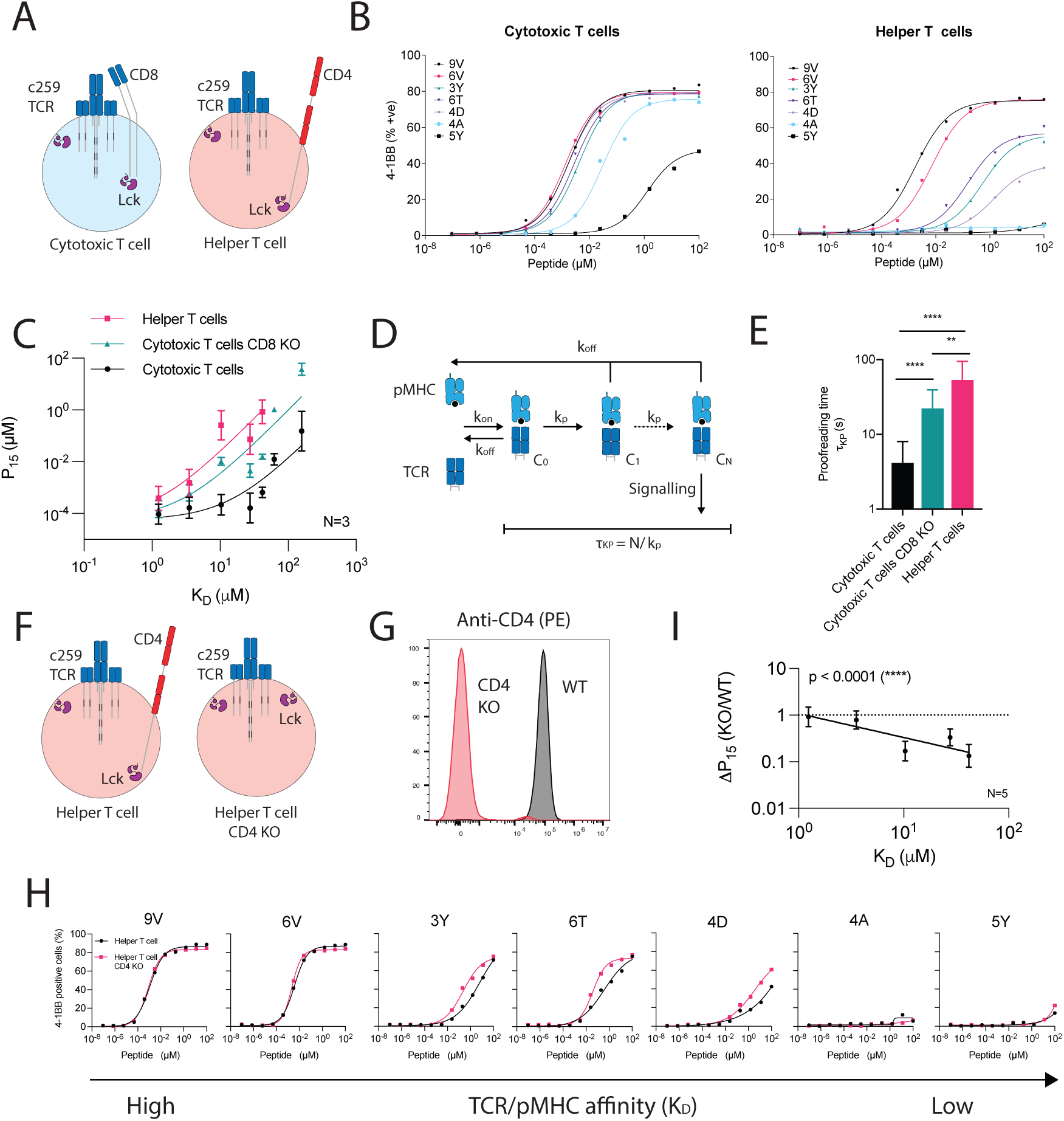
The CD4 co-receptor enhances the discrimination of helper cells expressing an MHC-I restricted TCR. **(A)** Schematic of helper and cytotoxic T cells transduced with the c259 TCR. **(B)** Representative ligand discrimination assays using helper and cytotoxic c259 TCR-T cells recognising peptides on U87 target cells. Expression of the 4-1BB activation marker was measured after a 20 hour co-culture. **(C)** Mean potency (P15) over TCR/pMHC affinity (K_D_) from N=3 independent blood donors (points) is fitted to the kinetic proofreading model (solid line). **(D)** Kinetic proofreading introduces a time-delay (*τ*_kp_) between pMHC binding (state *C*_0_) and TCR signalling (state *C_N_*) that selectively reduces signalling to low-affinity ligands. **(E)** Fitted time-delay from the data in panel (C). F-test compares the time-delay between conditions. **(F)** Schematic of helper WT and CD4 KO c259 TCR-T cells. **(G)** Flow cytometry staining of CD4 in WT and CD4 KO helper T cells. **(H)** U87 cells were titrated with each of the 7 NY-ESO-1 peptides to stimulate WT or CD4 KO helper c259 TCR-T cells. 4-1BB expression was measured after 20 hours. **(I)** Fold change in potency (P15) between CD4 KO and WT helper T cells from (H) is plotted over TCR/pMHC affinity (K_D_). Dashed line indicates fold change of 1. Significance of non-zero slope was assessed by an F-test Data in (B), (G) and (H) are representative of at least N=3 independent experiments with different blood donors. Data in (I) is shown as means ± SDs of N=5 independent experiments with different blood donors. ns not significant, *p*<*0.05, **p*<*0.01.

The degree to which T cells are able to respond to lower-affinity antigens is partly determined by a kinetic proofreading mechanism that introduces a time-delay between pMHC binding and TCR signalling (22, 29) (Fig. 3D). This time-delay is thought to be determined by biochemical steps that follow pMHC binding, including phosphorylation of ITAMs and ZAP70 by Lck, ZAP70 auto-phosphorylation, and the bridging of ZAP-70 and LAT by Lck (30–32). By fitting the proofreading model directly to the potency over pMHC affinity data (Fig. 3E), we confirmed that the time-delay for helper T cells is even larger than CD8 KO cytotoxic T cells. Thus, high levels of ligand discrimination for helper T cells cannot be explained simply by the absence of CD8 co-receptor alone.

Helper T cells express the CD4 co-receptor that like CD8 has an intracellular association with Lck, but unlike CD8 cannot bind the MHC-I antigens targeted by the c259 TCR. We hypothesised that the presence of the incompatible CD4 co-receptor could be responsible for the enhanced discrimination of helper T cells. Indeed, CD4 KO helper T cells displayed improved activation to lower affinity peptides, reducing ligand discrimination compared to wild-type helper T cells (Fig. 3F-I). Therefore, the incompatible CD4 co-receptor increases the ligand discrimination of helper T cells targeting pMHC-I antigens.

### Expression of the incompatible CD4 co-receptor in cytotoxic T cells enhances their ligand discrimination

Since the CD4 co-receptor increased the ability of helper T cells to discriminate ligands using an MHC-I restricted TCR, we examined whether it could also do this in cytotoxic T cells (Fig. 4A). Indeed, expression of CD4 in cytotoxic T cells selectively reduced activation and target killing against lower affinity pMHCs, without affecting responses to the high affinity cognate peptide (Fig. 4B, S6A). Moreover, expression of CD4 in CD8 KO cytotoxic T cells synergised to produce T cells with extremely high levels of discrimination (Fig. 4B-C, S6B-C). For example, whereas wild-type T cells can respond to the lower-affinity 4D peptide, these CD8→CD4 co-receptor switch T cells ignore this same antigen unless its concentration was increased by a dramatic ∼3000-fold. Thus, a CD8→CD4 co-receptor switch dramatically increased the ligand discrimination of cytotoxic T cells.

**Figure 4:**
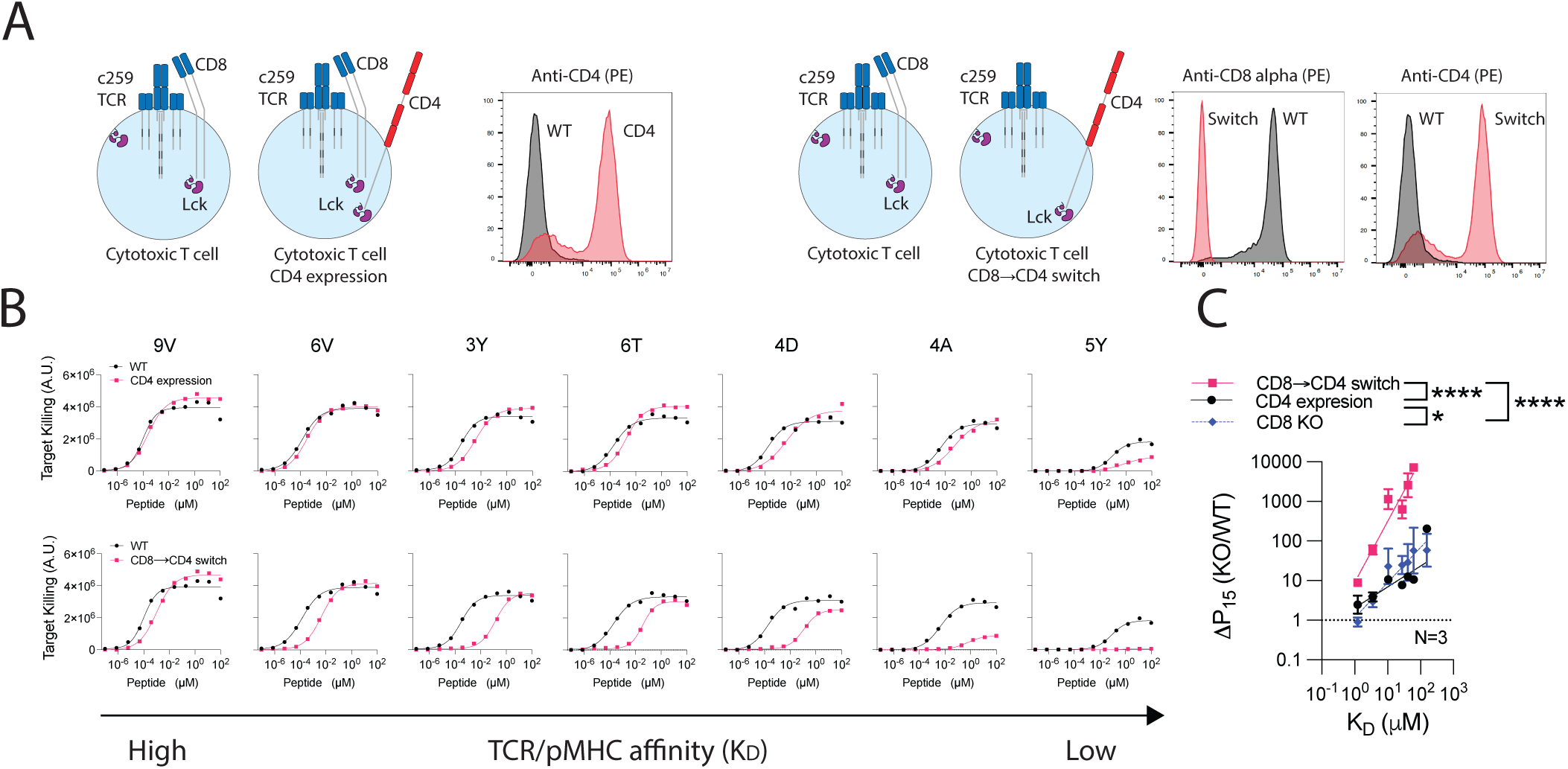
Expression of the incompatible CD4 co-receptor in cytotoxic T cells enhances ligand discrimination. **(A)** (Left) Schematic of CD4 expression in cytotoxic T cells and flow cytometry staining of CD4 expression. (Right) Schematic of CD8→CD4 co-receptor switch T cells and flow cytometry staining of CD4 and CD8 expression. **(B)** U87 cells were titrated with each of the 7 NY-ESO-1 peptides to stimulate (Top) WT or CD4 expressing cytotoxic T cells or (Bottom) WT or CD8→CD4 co-receptor switch cytotoxic T cells. Target killing was measured after 20 hours. **(C)** The fold change in Potency (P15) between the indicated modified and WT cytotoxic T cells over TCR/pMHC affinity (K_D_). Data for CD8 KO is shown from Fig 1. Data in (A) and (B) are representative of N=3 independent experiments with different blood donors. Data in (C) is shown as means ± SDs. P values were determined by F-test. *p*<*0.05, ****p*<*0.0001.

### CD8→CD4 co-receptor switch cytotoxic T cells display reduced cross-reactivity whilst maintaining potent target killing

We next used three methods to examine how the increase in ligand discrimination that we report impacts T cell cross-reactivity.

In a pooled peptide library that contains a random mixture of peptides, it is expected that the majority of peptides that bind the TCR would do so with low affinity. As a result, we predicted that increasing ligand discrimination would reduce T cell cross-reactivity to a pooled library (Fig. 5A). We stimulated T cells with target cells pulsed with a random pooled 9-mer peptide library, where each position can be any amino acid except cysteine, with a theoretical diversity of 19^9^ peptides. Cytotoxic T cells expressing the c259 TCR killed target cells pulsed with the random peptide mixture, but reduced cross-reactive killing was observed in CD8 KO and especially in CD8→CD4 co-receptor switch T cells (Fig. 5B).

**Figure 5:**
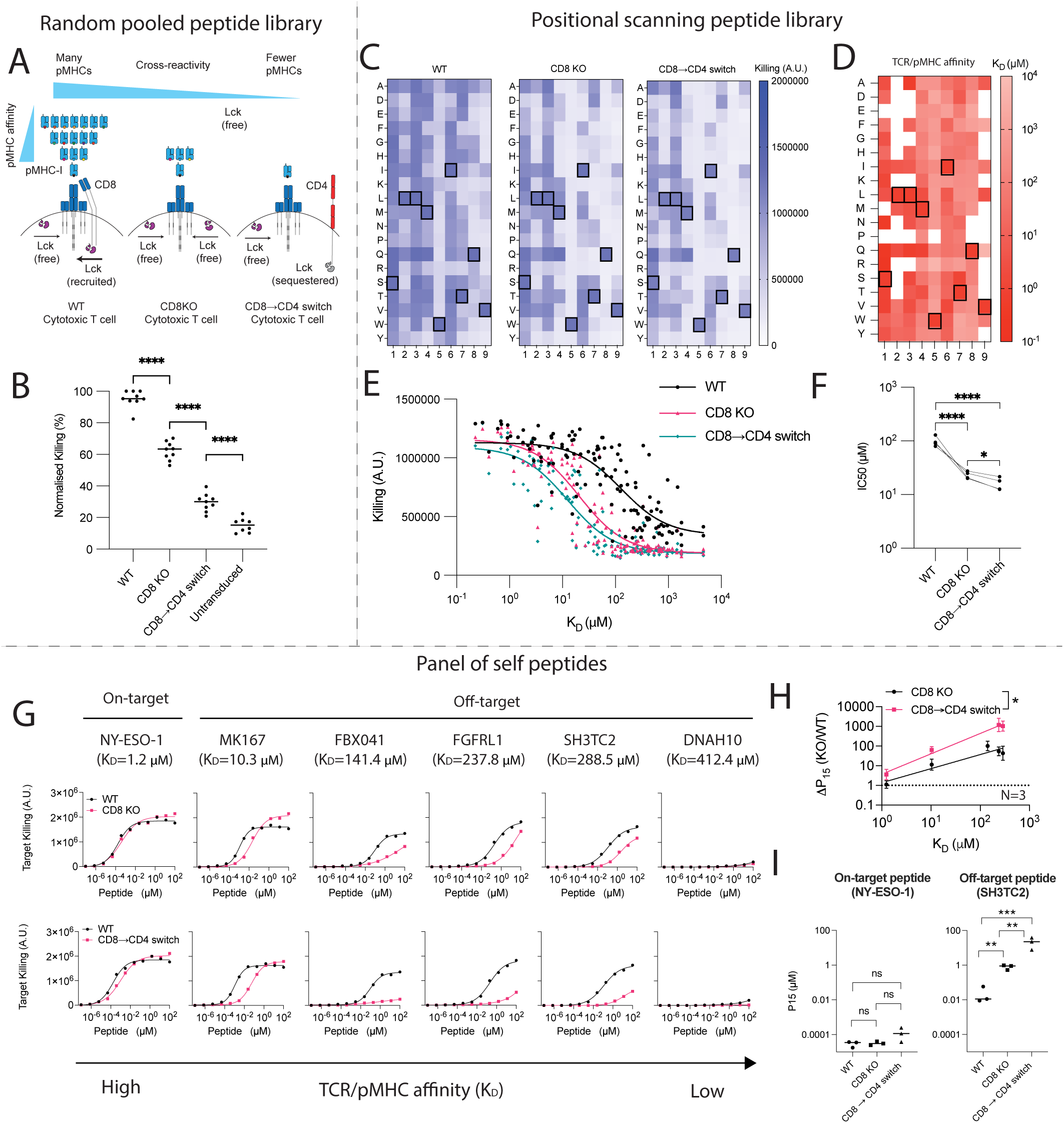
CD8→CD4 co-receptor switch cytotoxic T cells display reduced cross-reactivity to peptide libraries and self peptides without compromising on-target potency. **(A-B) Pooled peptide library. (A)** Schematic of the predicted cross-reactivity of WT, CD8 KO or CD8→CD4 co-receptor switch cytotoxic T cells. **(B)** U87 cells were loaded with a pooled 9-mer peptide library to stimulate WT, CD8 KO or CD8→CD4 co-receptor switch cytotoxic c259 TCR-T cells. Target killing was measured after 20 hours. Each data point represents a technical replicate from N=3 independent experiments with different blood donors. P values were determined by paired t-test. **(C-F) Positional Scanning Peptide Library. (C)** U87 cells were individually loaded with 0.1 µM of each of the 163 peptides in the positional library and co-cultured with T cells. Target killing was measured after 20 hours. Boxed amino acids represent the NY-ESO-1 peptide SLLMWITQV. **(D)** Affinity between c259 TCR and each pMHC in the positional library determined at 37 degrees by a high-throughput SPR method. Mean K_D_ values are shown from N=3 independent experiments. Boxed amino acids represent the NY-ESO-1 peptide SLLMWITQV. White boxes represent peptides without detectable MHC binding. **(E)** Target cell killing from (C) plotted over the TCR/pMHC K_D_ from (D). **(F)** IC50 from (E) is plotted with each data point representing an independent experiment with different blood donors. Data in (C) and (E) are representative data from N=4 independent experiments with different blood donors. P values were determined by paired t-test. **(G-I) Predicted self-peptides. (G)** U87 cells were titrated with each of the predicted self-peptides to stimulate (Top) WT or CD8 KO cytotoxic c259 TCR-T cells or (Bottom) WT or CD8→CD4 co-receptor switch cytotoxic c259 TCR-T cells. Target killing was measured after 20 hours. **(H)** Fold change in Potency (P15) between modified and WT T cells from (G) is plotted over TCR/pMHC affinity (K_D_). Data is shown as means ± SDs. P values were determined by F-test. **(I)** Potency (P15) from (G) is plotted for the indicated peptides. Each data point represents an independent experiment. P values were determined by paired t-test. Data in (G) are representative of N=3 independent experiments with different blood donors. ns not significant, *p*<*0.05, **p*<*0.01, ***p*<*0.001, ****p*<*0.0001.

A positional scanning library includes all single amino acid changes relative to a target peptide (163 NY-ESO-1 variant peptides in the present case). Although cytotoxic c259 TCR-T cells killed targets expressing many of these peptides, CD8 KO cells and CD8→CD4 co-receptor switch T cells display reduced killing to many of these peptides with the exception of the target peptide (Fig. 5C). To confirm that this reduced cross-reactivity was a result of increased ligand discrimination based on affinity, we developed a workflow to use a high-throughput SPR-based instrument to accurately and rapidly measure all 163 TCR/pMHC affinities (Fig. 5D, S7, Table S2). As predicted, the reduced cross-reactivity of CD8 KO and CD8→CD4 co-receptor switch T cells was dependent on affinity with reduced responses observed only to lower-affinity interactions (Fig. 5E-F).

Data from positional scanning libraries can also be used to predict TCR off-target cross-reactivities and this method has previously been used to predict potential self peptides recognised by the c259 TCR (8). We found that c259 TCR-T cells responded to a subset of these predicted peptides whose affinity we then measured by SPR (Fig. S8, Table S3). The CD8 KO and especially the CD8→CD4 co-receptor switch T cells displayed reduced responses to target cells presenting these cross-reactive self peptides (Fig. 5G-H, S9). Importantly, this reduced cross-reactivity did not compromise potency to the on-target NY-ESO-1 cancer antigen (Fig. 5I). Thus, co-receptor switching can reduce T cell cross-reactivity to increase the safety of TCR-T cell therapies.

## Discussion

It has been estimated that a single T cell can recognise over 10^6^ different peptides (33, 34). This cross-reactivity is an essential feature of adaptive immunity, enabling the limited number of T cell clones within an organism to provide protection against a much larger number of pathogenic peptides. However, T cell cross-reactivity poses a significant challenge to the success of TCR-T therapies as it can lead to lethal off-target toxicities. Identifying safe and effective TCRs remains a critical bottleneck in the development of new therapies. Despite this binding cross-reactivity, T cells use kinetic proofreading to discriminate between high and low affinity peptides (22, 29). Since ligand discrimination emerges not only from TCR binding but from TCR signalling (31, 32), we hypothesised that modifying T cell co-signalling receptors involved in this signalling pathway could be exploited to increase T cell ligand discrimination and reduce cross-reactivity without modifying the TCR. We have demonstrated the ability to increase and decrease ligand discrimination by genetic knockout and/or expression of the surface molecules CD5, CD4, and CD8 in helper/cytotoxic T cells. The CD8→CD4 co-receptor switch produced super selective T cells that display a striking increase in ligand discrimination and reduced cross-reactivity to a pooled and positional scanning libraries, and to self peptides without impacting on-target potency.

The CD8 co-receptor plays an essential role in thymic selection but its role in ligand discrimination is debated. Previous work established that CD8 increases T cell activation by stabilising the extracellular TCR-pMHC interaction (35) and by recruiting Lck to the signalling subunits of the TCR-CD3 complex (36). It has been proposed that CD8 can selectively stabilise high-affinity TCR/pMHC interactions through a positive feedback that amplifies differences in binding affinity and hence enhances ligand discrimination (35, 37). On the other hand, it has been suggested that CD8 slows the dissociation rate of TCR/pMHC interactions (38), which preferentially increases the sensitivity to low affinity peptides and hence reduces ligand discrimination (39–41). Our systematic analyses support the latter hypothesis, showing that CD8 KO selectively reduces activation towards lower-affinity antigens and hence CD8 KO increases ligand discrimination.

We found that CD4+ helper T cells display higher levels of ligand discrimination compared to CD8+ cytotoxic or CD8 KO cytotoxic T cells recognizing MHC-I antigens. This suggested that the CD4 co-receptor, which binds MHC-II, might further increase ligand discrimination. We confirmed this by showing that CD4 KO in helper T cells reduced their ligand discrimination and expression of CD4 in WT or CD8 KO cytotoxic T cells enhanced their ligand discrimination. These findings are consistent with the Lck sequestration model first proposed to understand thymocyte development. This model postulates that CD4/CD8 co-receptors inhibit signalling when they are not able to recognise the ligand recognised by the TCR, by sequestering Lck from the TCR (42–44). We suggest that removal of a compatible co-receptor or the introduction of an incompatible co-receptor increases the proofreading time-delay between pMHC binding and TCR signalling leading to enhanced ligand discrimination (Fig. 3).

Whilst increasing the discrimination of therapeutic TCRs can increase their safety, decreasing ligand discrimination has been proposed as an attractive strategy to increase activation against lower affinity immune escape peptide variants in tumours with high genomic instability (45). We have identified CD5 KO as a candidate modification to decrease T cell ligand discrimination and our findings are consistent with its negative regulatory function that fine-tunes TCR signalling to maintain T cell tolerance and reduce the risk of autoimmunity (24, 46). Although reducing the function of CD5 has been shown to enhance anti-tumour activity in TCR-T and CAR-T cells (47–49), this may be a double-edge sword because it would also increase cross-reactivity and hence the risk of autoimmune toxicities. Similarly, on-going clinical trials have engineered CD4+ helper T cells to express the CD8 co-receptor to increase their potency (50) but our results suggest that this may increase their cross-reactivity and the risk of autoimmune toxicities.

Overall, we have demonstrated that super selective T cells with reduced cross-reactivity and enhanced ligand discrimination can be generated without impacting on-target potency and importantly, without modifying the TCR. We have applied the method to the clinical a3a and c259 TCRs showing that it can abolish functional cross-reactivity to self peptides. A limitation of this method is that if a TCR does not bind its target cancer peptide with high affinity, its potency may be reduced by co-receptor switching. Therefore, affinity-maturation might be required for lower-affinity therapeutic TCRs. Given that these super selective T cells are generated by modifying genes extrinsic to the TCR, it has the potential to dramatically increase the safety of TCR-T cell therapies regardless of the therapeutic TCR that is used.

## Funding

The work was funded by a Wellcome Trust Senior Fellowship in Basic Biomedical Sciences (207537/Z/17/Z to OD) and by UKRI-Biotechnology and Biological Sciences Research Council (BB/T008784/1 to JCC).

## Open access

This research was funded in whole, or in part, by the Wellcome Trust [207537/Z/17/Z]. For the purpose of Open Access, the author has applied a CC BY public copyright licence to any Author Accepted Manuscript version arising from this submission.

## Author contributions

Conceptualization (JCC, OD), Data Curation (JCC, AH, MK, AS), Formal Analysis (JCC, AH), Funding Acquisition (OD), Investigation (JCC, AH, MK, AS), Methodology (JCC, VA, AH, MK, AS, PAvDM, OD), Project Administration (OD), Supervision (PAvDM, OD), Visualization (JCC), Writing – Original Draft (JCC, OD), Writing – Review & Editing (JCC, AH, MK, AS, PAvDM, OD)

## Materials & Methods

### Cell culture

U87 and HEK cell lines were cultured at 37°C and 10% CO2 in DMEM D6429 media (Sigma-Aldrich) supplemented with 10% FBS, 50 µg/mL Streptomycin and 50 units/mL Penicillin.

T2 cells and Nalm6 cells were cultured at 37°C and 10% CO2 in RPMI 1640 (Sigma-Aldrich) supplemented with 10% FBS, 50 µg/mL Streptomycin, 50 units/mL Penicillin.

Primary human T cells were isolated from leukocyte cones and cultured at 37°C and 10% CO2 in RPMI 1640 (Sigma-Aldrich) supplemented with 10% FBS, 50 µg/mL Streptomycin, 50 units/mL Penicillin and 50 U/mL IL2.

### Lentivirus production

0.8 Million HEK 293T cells were seeded in a 6-well plate (Day 1) and incubated overnight. Cells in each well were co-transfected (Day 2) using X-tremeGENE™ HP (Roche) with 0.8 µg of the appropriate lentiviral transfer plasmid encoding an antigen receptor (1G4 TCR or c259 TCR) and the lentiviral packaging plasmids: pRSV-Rev (0.25 µg), pMDLg/pRRE (0.53 µg), and pVSV-G (0.35 µg). The media was replaced 18 hours following transfection (Day 3). 24 hours after the media exchange, the supernatant from one well was harvested, filtered and used for the transduction of 1 Million human T cells (Day 4).

### Production of TCR transduced primary human T cells

T cells were isolated from anonymised leukocyte cones (Day 3) purchased from the NHS Blood Donor Centre at the John Radcliffe Hospital (Oxford University Hospitals). As a result of the anonymised nature of the cones, biological sex and gender were not variables in the present study and were therefore randomised, and as a result the authors were blinded to these variables. RosetteSep™ Human CD8+ Enrichment Cocktail (STEMCELL Technologies) was used for cytotoxic T cells or CD4+ T Cell Enrichment Cocktail (STEM-CELL Technologies) for helper T cells. The enrichment cocktail was added at 150 µl/mL of sample and incubated at RT for 20 minutes. The sample was diluted with an equal volume of PBS and layered on Ficoll® Paque Plus (Cytiva) density gradient medium at a 0.8:1 ratio (Ficoll®:Sample).

The sample was centrifuged at 1200 g for 30 minutes (brake off). Cells at the interface of the Ficoll® media and plasma were collected (Buffy coat) and washed twice (Centrifuged at 500 g for 5 minutes). Cells were resuspended in complete RPMI media supplemented with IL2 (50 U/mL) at a density of 1 Million cells per mL. Dynabeads® Human T-Activator CD3/CD28 (Thermofisher) were added (1 Million beads per mL) and cells were incubated overnight.

1 Million cells were transduced with the filtered lentiviral supernatant (Day 4). On Day 6 and on Day 8, 1 mL of media was removed and replaced with 1 mL of fresh medium. On Day 9, Dynabeads® were removed using a magnetic stand (6 days following isolation). Cells were resuspended in fresh media every other day at a density of 1 Million per mL and used for co-culture experiments. 17 days following isolation T cells were discarded.

### CRISPR/Cas9 knock-out of T cell proteins

Cas9 ribonucleoproteins (RNPs) were prepared by mixing 8.5 µg of TruCut Cas9 protein v2 (Thermofisher) with 150 pmol of sgRNA mix (Truguide synthetic grna, Thermofisher) and Opti-MEM (Gibco) to a final volume of 5 µl. The RNPs were incubated for 15 minutes at room temperature.

1 Million freshly isolated T cells were washed with Opti-MEM (Gibco) and re-suspended at a density of 20 Million per mL. The T cells were mixed with the RNPs and transferred into a BTX Cuvette Plus electroporation cuvette (2mm gap, Harvard Bioscience). The cells were electroporated using a BTX ECM 830 Square Wave Electroporation System (Harvard Bioscience) at 300 V, 2 ms. Immediately following electroporation, the cells were transferred to complete RPMI media supplemented with IL2 and Dynabeads® Human T-Activator CD3/CD28 (Thermofisher) were added.

The following sgRNA sequences were used:

#### CD8 alpha knock-out

Guide 1: ATACTGTTGTGCGCACATCG

Guide 2: GTTAGACGTATCTCGCCGAA

Guide 3: GCTGCTGTCCAACCCGACGT

Guide 4: GAGCAAGGCGGTCACTGGTA

#### CD5 knock-out

Guide 1: GCAGACTTTTGACGCTTGAC

Guide 2: CCGTTCCAACTCGAAGTGCC

Guide 3: ATCATCTGCTACGGACAACT

Guide 4: AGGTCTACCTCAAGGACGGA

#### CD43 knock-out

Guide 1: GGCTCGCTAGTAGAGACCAA

Guide 2: GCACCAATGGAAGTCCAAAG

Guide 3: AGGTTGTTGGCTCAGGTAAA

#### CD2 knock-out

Guide 1: CAAGGCACCCCAGGTTTCCA

Guide 2: CAAAGAGATTACGAATGCCT

Guide 3: CTTGTAGATATCCTGATCAT

Guide 4: GCATCTGAAGACCGATGATC

#### CD11a knock-out (LFA-1)

Guide 1: CTTTGGATACCGCGTCCTGC

Guide 2: CAAGTACTTGGAGGTATAGT

Guide 3: GTAACACAGGCCACTCAGAT

Guide 4: GUAGCUCGAGGCCGGCGCUG

#### CD4 knock-out

Guide 1: GTCAGCGCGATCATTCAGCT

Guide 2: GAGGTGCAATTGCTAGTGTT

Guide 3: AACTGTAAAGGCGAGTGGGA

Guide 4: CTGTTTTCGCTTCAAGGGCC

### Negative selection of T cell knock-out cells

T cells with residual target protein expression were depleted by antibody staining and bead pull-down. T cells were re-suspended in MACS Buffer (PBS, 0.5% BSA, 2 mM EDTA) at a density of 100 Million cells per mL. Cells were stained with 5 µl of the corresponding PE-labelled antibody per million cells for 15 minutes at 4°C, washed with MACS and re-suspended at a density of 100 Million cells per mL. 1 µl of MojoSort anti-PE nanobeads (Biolegend) were added per million cells and incubated on ice for 15 minutes. The cells were washed with MACS and the beads were pulled-down magnetically. The supernatant containing the negatively selected cells was collected.

### Cellular co-culture assays

50 000 U87 cells in 100 µl of DMEM were seeded per well in a 96-well Flat-bottom plate and incubated overnight. Alternatively, 100 000 T2 cells were placed in each well of a 96-well Flat Bottom plate. Peptides were diluted in DMEM to the appropriate concentration, added to each well containing cells and incubated for 60 minutes at 37°C, 10% CO2. The media was discarded and 50 000 T cells were added to each well in 200 µl of RPMI media. Cells were incubated for 20 hours at 37°C, 5% CO2. Supernatants were collected for cytotoxicity and ELISA analysis. 25 µl of 100 mM EDTA PBS were added to each well containing the cells and samples were incubated for 5 minutes at 37°C, 5% CO2. Cells were detached by thoroughly pipetting each well and transferred to a 96-well V-bottom plate.

### Flow Cytometry

Cells were stained for 20 minutes at 4°C, washed with PBS and analysed using a BD X-20 flow cytometer or Cytoflex LX Flow cytometer (Beckman Couter). The starting cell population was gated on a linear SSC-A/FSC-A plot. Single cells were discriminated on a linear FSC-H/FSC-W plot. In co-culture experiments using U87 cells, T cells were gated as CD45 positive. In co-culture experiments using Nalm6 or T2 cells, T cells were gated as CD3 positive. Positive/negative populations were determined with negative controls. Data was analysed using FlowJo v10, RRID:SCR008520 (BD Biosciences) and GraphPad Prism, RRID:SCR002798 (GraphPad Software).

### Cytotoxicity assay

Target cell lines were engineered to express the Nluc luciferase (51). A Coelenterazine (CTZ) 2 mM stock solution was prepared in methanol, aliquoted and stored at -80°C. Supernatant from co-culture assays was mixed in a 1:1 ratio with PBS 10 µM CTZ and luminescence was read using a SpectraMax M3 microplate reader (Molecular Devices).

### Cytokine ELISA

Invitrogen Human IFN*γ* ELISA kits (Thermo Fisher Scientific) were used following the manufacturer’s protocol to quantify levels of cytokine in diluted T cell supernatant. A SpectraMax M3 microplate reader (Molecular Devices) was used to measure absorbance at 450 nm and 570 nm.

### Surface Plasmon Resonance

All SPR experiments were carried out in the Dunn School SPR facility following the methods published on (22). Briefly, c259 TCR/pMHC steady-state binding affinities were measured on a Biacore T200 (GE Healthcare) with a CAP chip using HBS-EP as running buffer. The CAP chip was saturated with streptavidin and biotinylated pMHCs were immobilised to the desired level. A titration of the TCR was flowed through at 37°C. CD58 was immobilised on a reference flow cell at matching levels to those of pMHCs on the remaining flow cells. The signal from the reference flow cell was subtracted (Single referencing) and the average signal from the closest buffer injection was subtracted (Double referencing). Steady-state binding affinity was calculated by fitting the one site-specific binding model (Response = B_max_ [TCR]/(K_D_ + [TCR]) on GraphPad Prism to double-referenced equilibrium RU values. The B*_max_* was constrained to the inferred B*_max_* from the empirical standard curve, relating maximal antibody binding to maximal TCR binding.

### Pooled Peptide Libraries

50 000 U87 cells in 100 µl of DMEM were seeded per well in a 96-well Flat-bottom plate and incubated overnight. The 9-mer pooled peptide library was diluted in DMEM to 100 µM, added to each well containing cells and incubated for 60 minutes at 37°C, 10% CO2. 50 000 T cells were added to each well in 200 µl of RPMI media. Cells were incubated for 20 hours at 37°C, 5% CO2. Supernatants were collected for cytotoxicity analysis.

### Positional Scanning Peptide Library SPR

To prepare pMHC complexes presenting the local peptide library, a disulfide-stabilized variant of the human MHC-I protein HLA-A*02:01 (DS-A2) was used (52). The DS-A2 protein was produced as described previously (52). Briefly, the DS-A2 and β2-microglobulin (β2m) subunits were produced in E. coli as inclusion bodies and solubilized in 8 M urea. The protein was then refolded in the presence of GlyLeu, a dipeptide that binds with low affinity to the peptide-binding cleft. The refolded DS-A2–β2m complexes were purified by size exclusion chromatography on a Superdex S75 10/300 column (GE Healthcare/Cytiva) in HBS-EP buffer (10 mM HEPES pH 7.4, 150 mM NaCl, 3 mM EDTA, and 0.05% Tween 20). Local-library peptides were loaded by incubating the DS-A2–β2m complex with each peptide for 2 h at room temperature. The pMHC complexes were stored at 4°C until use within 24 h.

Soluble c259 TCR was produced as separate TCRα and TCRβ chains in *E. coli*. Both chains were recovered as inclusion bodies, solubilised in 100 mM Tris-HCl (pH 8.0), 8 M urea, 2 mM DTT and stored in aliquots at -70°C. For refolding, 30 mg of each TCR chain was added to 1 L of refolding buffer (150 mM Tris-HCl (pH 8.0) 3 M urea, 200 mM Arg-HCl, 0.5 mM EDTA, 0.1 mM PMSF) and stirred for 1 h at 4°C. This was followed by dialysis in 10 L 10 mM Tris-HCl (pH 8.5) buffer for 3 days in total, with the dialysis buffer changed after 1 day. The refolded c259 TCR was purified using anion exchange chromatography (HiTrap Q HP; Cytiva), followed by size exclusion chromatography (Superdex 200 Increase; Cytiva) in HBS-EP Buffer. Purified c259 was used within 48 h.

High-throughput affinity measurements of c259 TCR binding to MHC loaded with the peptide library were performed using LSA or LSA^XT^ (Carterra). Each pMHC was immobilised via biotin-streptavidin binding on a different spot of the SAHC30M biosensor (Carterra) for 20 min, resulting in immobilisation levels between 200 and 900 RUs. Measurements were performed in HBS-EP Buffer at 37 °C. A 2-fold dilution series of c259 TCR was prepared in HBS-EP buffer, with the highest concentration between 100 - 130 µM. Starting with the highest dilution, increasing concentrations of c259 were injected over the chip for 5 min, followed by 5 -10 min of dissociation, without regeneration. Afterwards, a *β*2m specific antibody (clone B2M-01 (Thermo Fisher Scientific) or BBM.1 (Absolute Antibody)) was injected for 10 min. The resulting data was analysed using Kinetics Software (Carterra). Any spikes were removed from the data, before referencing against empty control spots or spots immobilised with CD86 at matching immobilisation levels. The final in a series 6 buffer injection before TCR injection was subtracted from the data for double referencing. Subsequently, the steady state binding RU was calculated by taking the average RU from over 20 seconds. Steady-state analysis was performed to obtain the K_D_ values. First, steady-state data was fitted with a one site-specific binding model (Response = B_max_ [TCR]/(K_D_ + [TCR]), with K_D_ and B_max_ unconstrained. We then constructed an empirical standard curve using high affinity pMHCs (K_D_ *<* 20µM) to relate maximal anti-*β*2m binding to TCR B_max_. Next, steady state data for all pMHCs were fitted with a one site-specific binding model with B_max_ constrained to the B_max_ inferred from the empirical standard curve. We excluded K_D_ values for peptides, where we observed little or no anti-*β*2m binding responses, indicating that the pMHC complex was unstable and lost the peptide over time (indicated as N/A in Table 2). We further excluded K_D_ values for pMHC that produced a TCR binding response of less than 5 RU (indicated as non-binders (NB) in Table 2).

### Data analysis

EC_50_ is calculated as the concentration of antigen required to elicit 50% of the maximum response determined for each condition individually whereas P_15_ is calculated as the concentration of antigen required to elicit 15% of the maximum activation.

The study is largely focused on comparing antigen sensitivity using EC_50_ or P_15_ measures, which we have found displays standard deviations of 0.2 (on log-transformed values). The smallest effective size that we aimed to resolve was 3-fold changes (a difference of 0.47 on log-transformed values) and a power calculation shows that this can be be resolved with a power of 80% (alpha at 0.05) using three samples in each group. Therefore, all experiments relied on a minimum of 3 independent donors.

## Data Availability

This study includes no data deposited in external repositories.

## Disclosure and competing interests statement

JCC, PAvdM, and OD have financial interests in a filed patent application related to this technology.

## Acknowledgements

We thank Malcolm Sim, Yardena Samuels, Shira Sagie-Groher, and Tomer Babu for helpful discussions. We thank Jonathan Popplewell for assisting with the LSA SPR experiments and Carterra Ltd for providing sensor chips.

## Ethics

Human research participants: Ethical approval was provided by the Medical Sciences Inter-divisional Research Ethics Committee (IDREC) at the University of Oxford (R51997/RE001).

## Supplementary Figures

**Figure S1:**
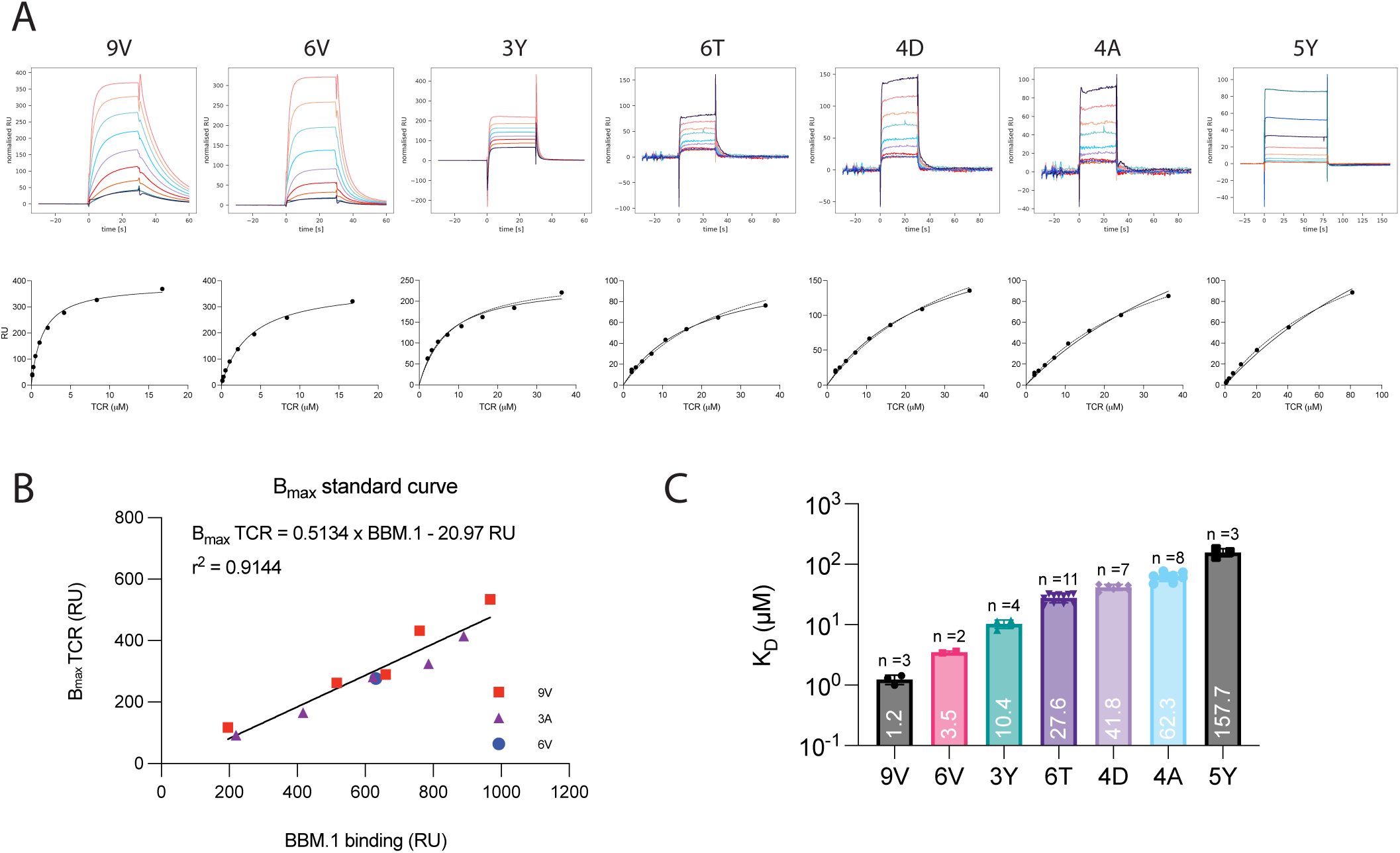
Establishing a panel of peptides that bind the c259 TCR with a range of affinities as measured by SPR at 37°C. **(A)** (Top) Representative SPR sensograms depicting injections of increasing concentrations of the c259 TCR. (Bottom) Representative steady-state curves of c259 TCR binding to different pMHCs. 3D affinity (K_D_) was calculated by constraining Bmax (dashed line) or fitting Bmax (solid line). **(B)** Empirical standard curve relating the binding of the BBM.1 antibody (x-axis) to the fitted TCR Bmax. Only data for the higher-affinity pMHCs is used to generate the standard curve. **(C)** Steady-state binding affinity for the selected 7-peptide panel. Barplot represents mean K_D_ ± SDs. The affinities were calculated by constraining Bmax to the value obtained from the standard curve in (B) based on the amount of BBM.1 antibody that bound the chip surface (see Methods for details). All data fitting was performed using a one site-specific binding model in GraphPad Prism.

**Figure S2:**
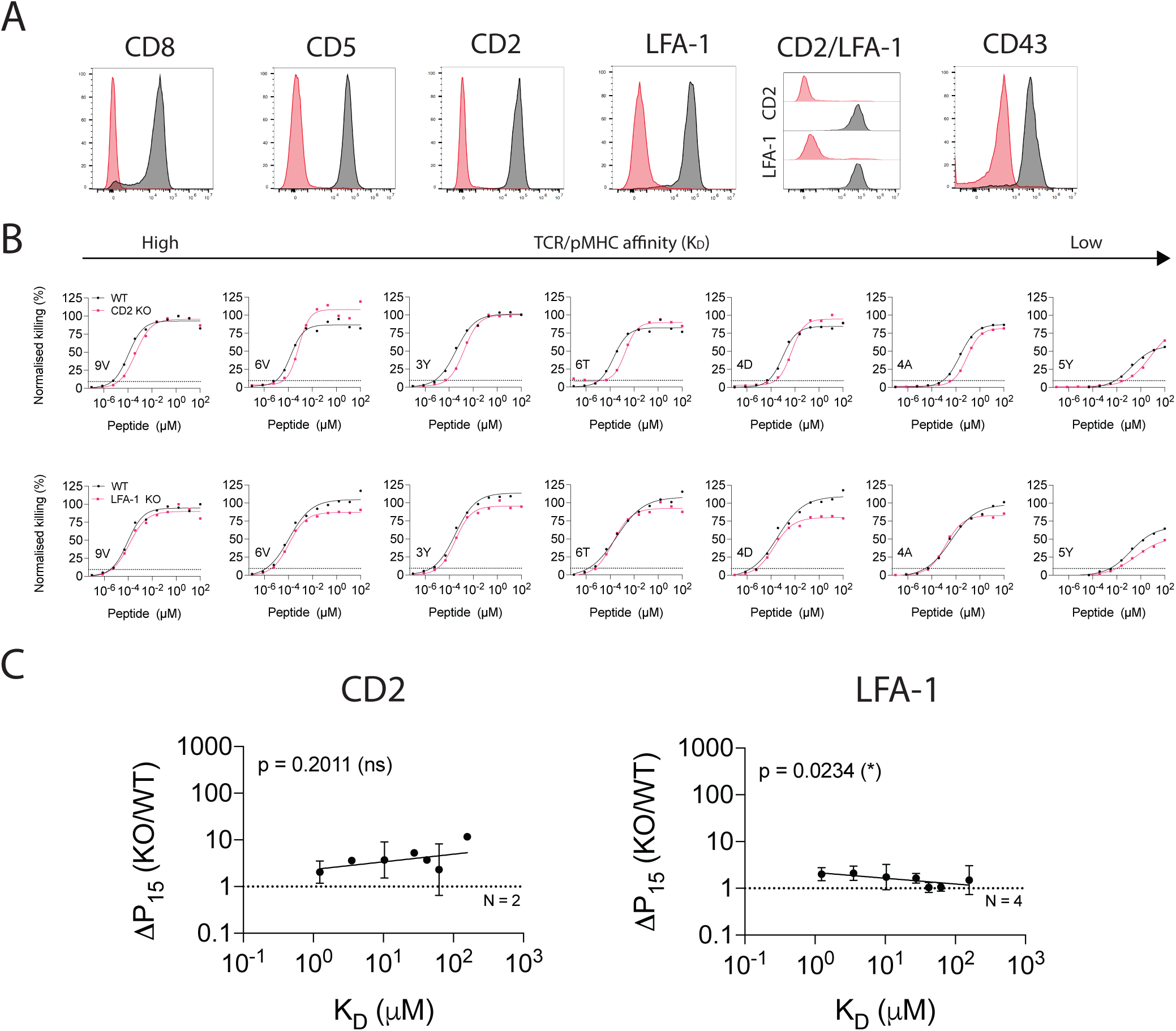
The impact of different T cell co-signalling receptors on ligand sensitivity and discrimination using target cell killing. **(A)** Flow cytometry staining of WT cells (Black) or KO T cells (Red). **(B)** U87 cells were titrated with each of the 7 NY-ESO-1 peptides to stimulate WT or KO c259 TCR-T cells. Killing of the target U87 cells was measured after 20 hours. Dashed line indicates potency (P15). **(C)** Change in potency over affinity as described in Fig. 1D. Data in (A) and (B) are representative of at least N=2 independent experiments with different blood donors. Dashed line in (C) indicates fold change of 1. Data in (C) is shown as means ± SDs. Significance of non-zero slope was assessed by an F-test.

**Figure S3:**
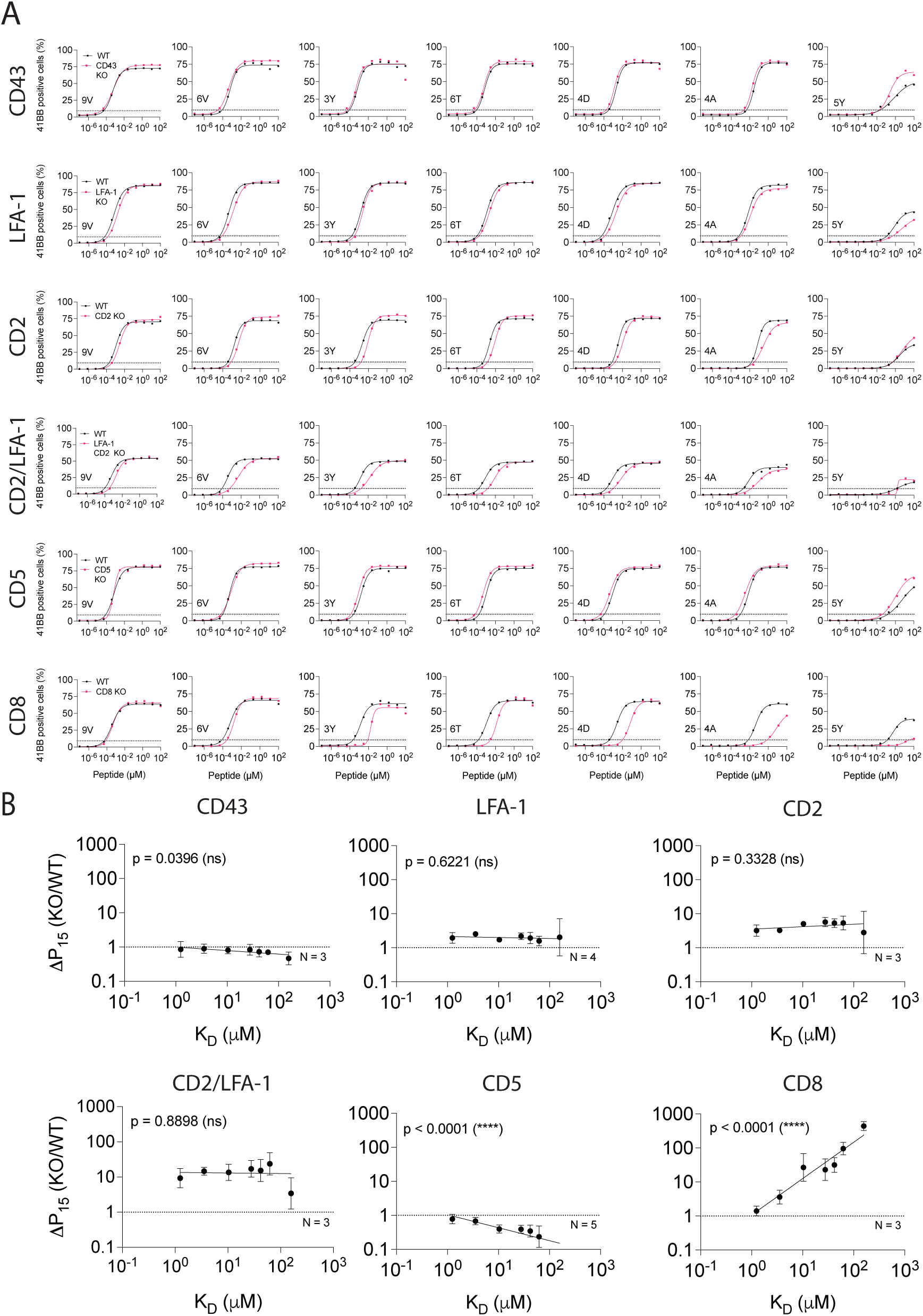
The impact of different T cell co-signalling receptors on ligand sensitivity and discrimination using 4-1BB activation marker. **(A)** Representative dose-response and **(B)** Change in potency over affinity as described in Fig. 1D for target killing. Data in (A) are representative of at least N=3 independent experiments with different blood donors. Dashed line in (B) indicates fold change of 1. Data is shown as means ± SDs. Significance of non-zero slope was assessed by an F-test.

**Figure S4:**
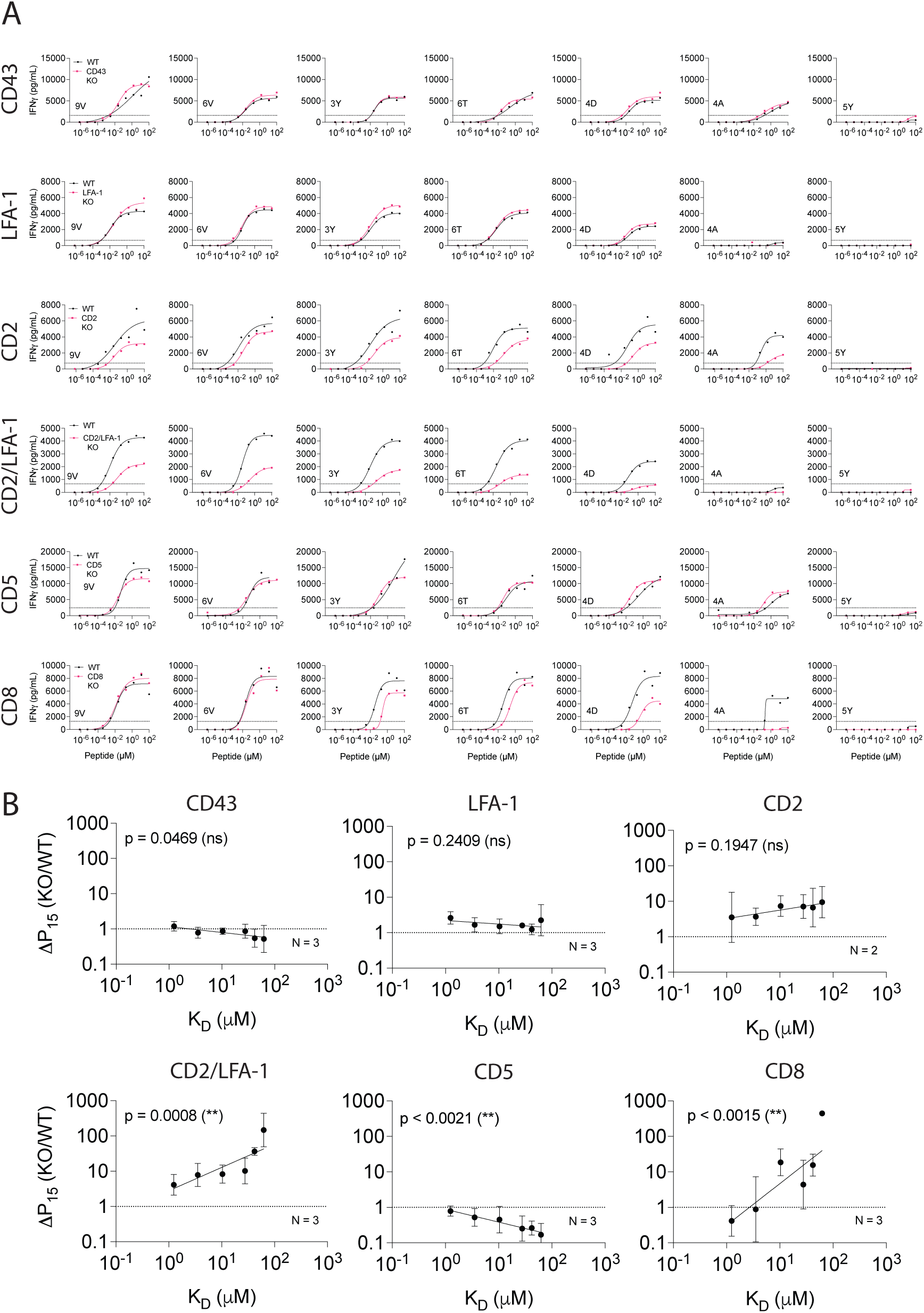
The impact of different T cell co-signalling receptors on ligand sensitivity and discrimination using the secreted cytokine IFN *γ*. **(A)** Representative dose-response and **(B)** Change in potency over affinity as described in Fig. 1D for target killing. Data in (A) are representative of at least N=2 independent experiments with different blood donors. Dashed line in (B) indicates fold change of 1. Data is shown as means ± SDs. Significance of non-zero slope was assessed by an F-test.

**Figure S5:**
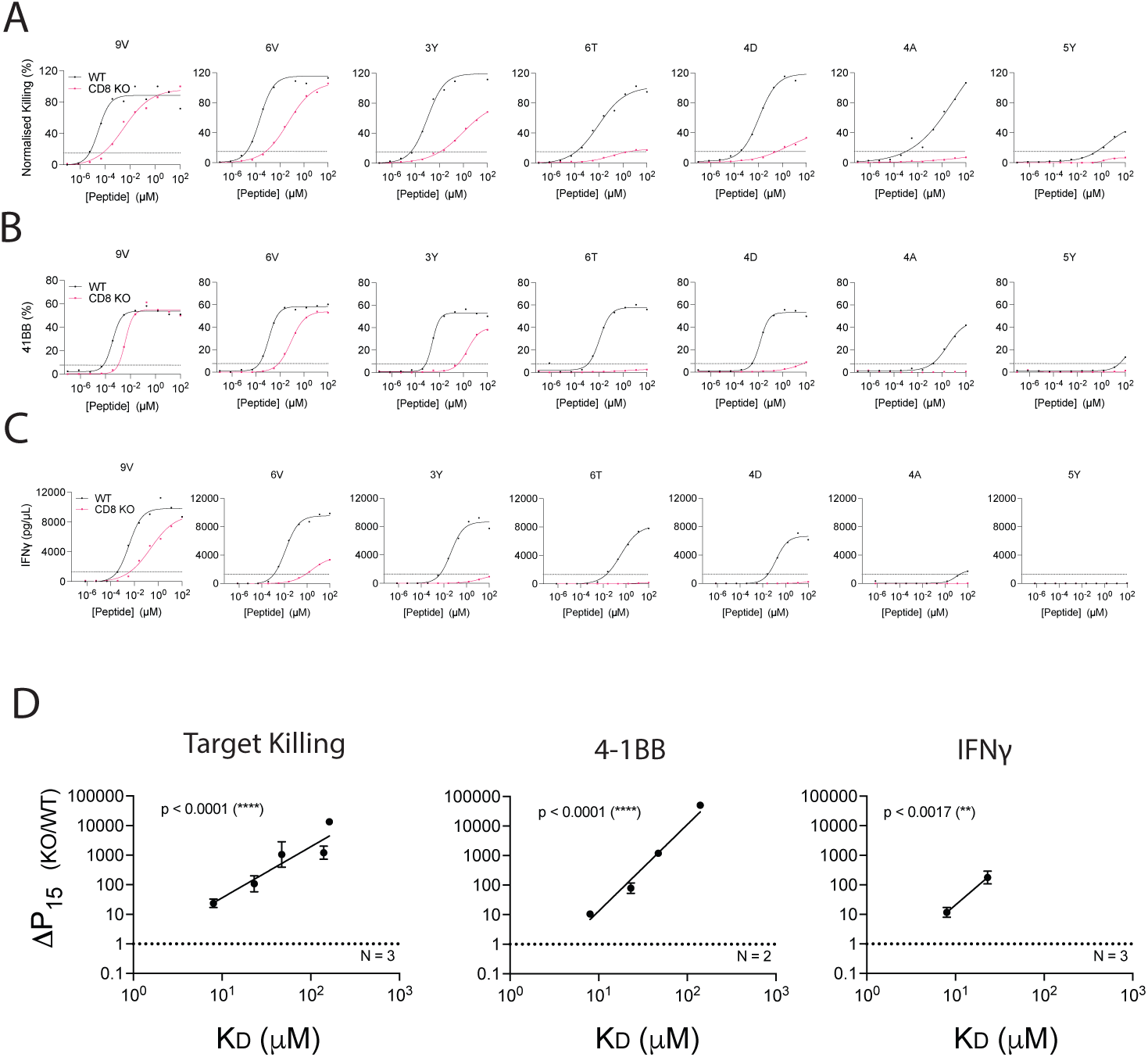
CD8 KO increases the ligand discrimination of the 1G4 TCR. **(A to C)** U87 cells were titrated with each of the 7 NY-ESO-1 peptides to stimulate WT or KO 1G4 TCR-T cells. **(A)** 4-1BB expression was measured after 20 hours. **(B)** Killing of the target U87 cells was measured after 20 hours. **(C)** IFN *γ* secretion was measured after 20 hours. **(D)** Fold change in potency (P15) between KO and WT T cells plotted over TCR/pMHC affinity (K_D_) (22). Dashed line indicates fold change of 1. Data is shown as means ± SDs. Data in (A), (B) and (C) are representative of at least N=2 independent experiments with different blood donors. Significance of non-zero slope was assessed by an F-test.

**Figure S6:**
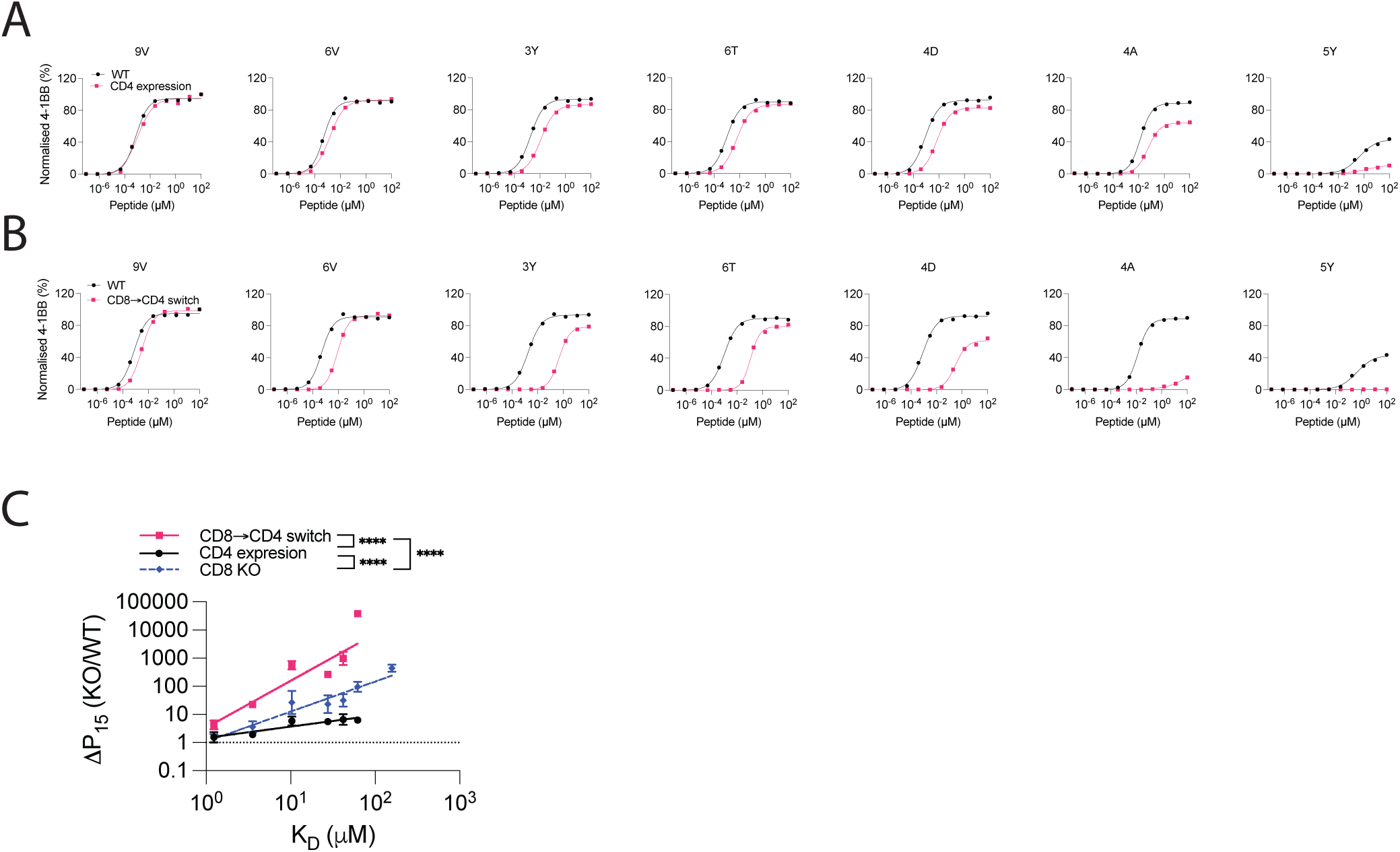
Expression of the incompatible CD4 co-receptor in cytotoxic T cells enhances ligand discrimination (4-1BB). **(A)** U87 cells were titrated with each of the 7 NY-ESO-1 peptides to stimulate WT or CD4 expressing cytotoxic c259 TCR-T cells. 4-1BB expression was measured after 20 hours. **(B)** U87 cells were titrated with each of the 7 NY-ESO-1 peptides to stimulate WT or CD8→CD4 co-receptor switch cytotoxic c259 TCR-T cells. 4-1BB expression was measured after 20 hours. **(C)** Fold change in potency (P15) between modified and WT T cells from **(A,B)** is plotted over TCR/pMHC affinity (K_D_). Data for CD8 KO is shown from Fig S3. Data is shown as means ± SDs. Data in (A) and (B) are representative of N=3 independent experiments with different blood donors. P value was determined by an F-test. ****p*<*0.0001.

**Figure S7:**
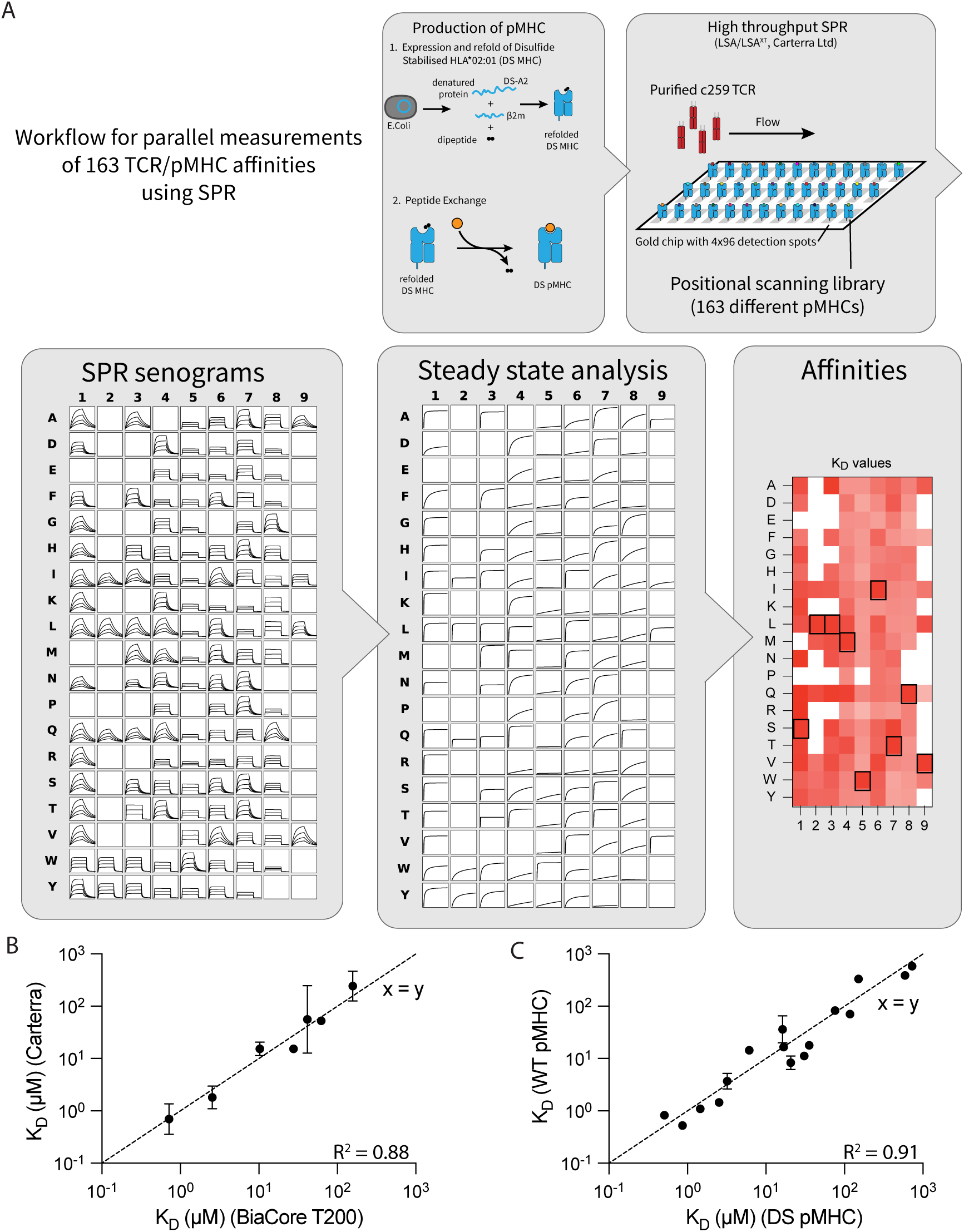
High-throughput measurements of c259 TCR affinities with the 163 pMHCs from the positional scanning library by SPR at 37°C. **(A)** Schematic of high-throughput SPR workflow. Step 1: Production of pMHCs presenting peptides from the positional scanning peptide library. Disulfide-stabilised HLA-A*02:01 (DS-A2) and *β*2m are expressed in *E.coli* as denatured protein chains, then refolded with a dipeptide. The dipeptide is exchanged with a peptide from the positional scanning peptide library by incubation. Step 2: High-throughput SPR setup. Using the LSA or LSA^XT^ instrument (Carterra) a pMHC carrying a each peptide from the library is immobilised in a separate detection spot on the chip. Soluble TCR is injected and flows over the entire chip. Step 3: Acquisition of SPR sensograms. Each detection spot simultaneously measures TCR binding over time for each peptide from the peptide library. Step 4: Calculation of affinity values. The steady-state binding response is plotted over TCR concentration to calculate K_D_ values using the constrained Bmax methods optimised for measuring ultra-low TCR/pMHC affinities (22). Step 5: The mean K_D_ values as heat map. **(B)** The K_D_ determined using the Carterra LSA/LSA^XT^ instruments agrees favourably with the K_D_ values determined using a standard BIAcore (T200). **(C)** The K_D_ determined using the disulfide-stablised MHC agrees favourably with the K_D_ determined using wild-type MHC for different peptides that bind the c259 TCR with a wide range of affinities.

**Figure S8:**
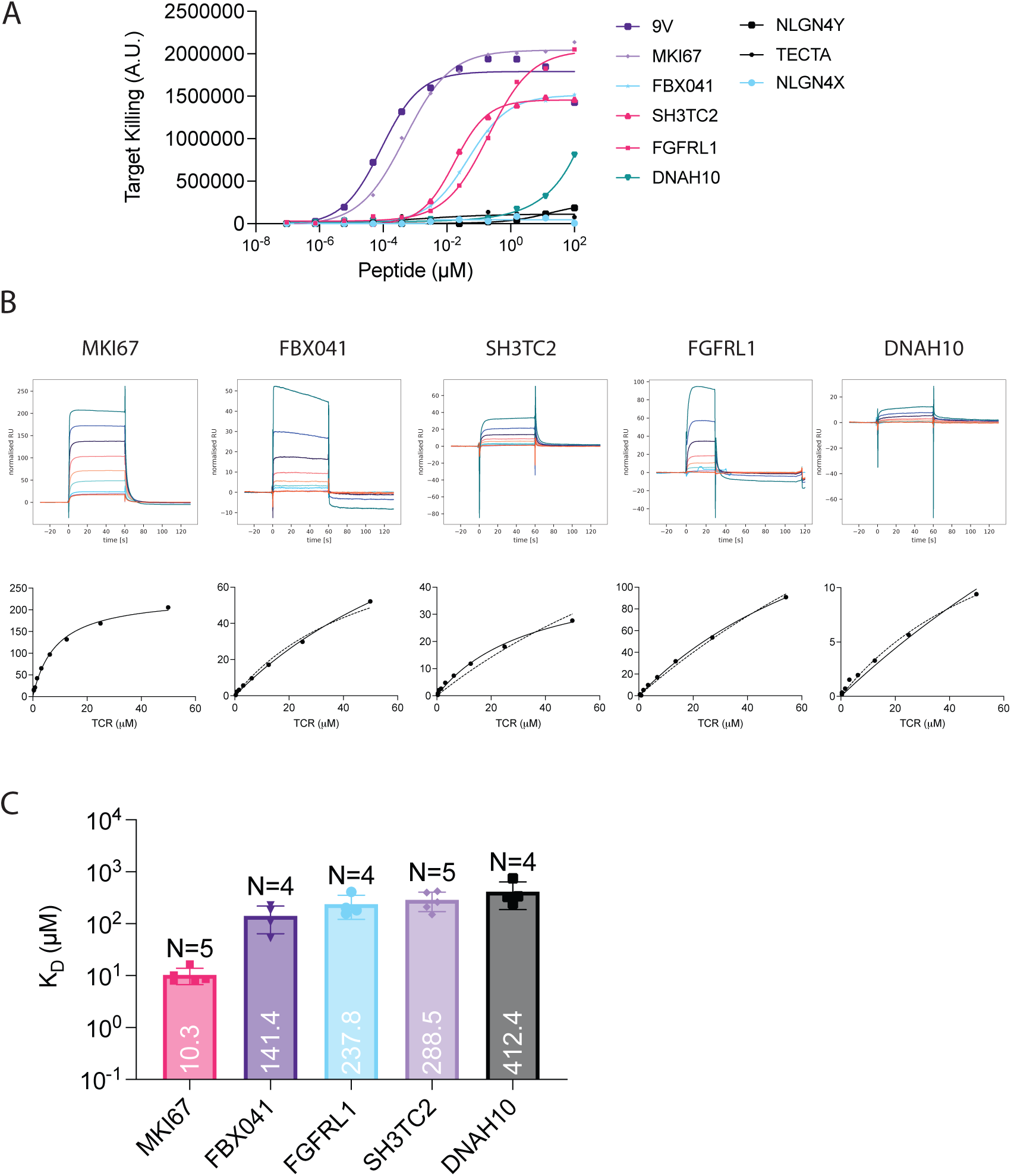
The c259 TCR affinity with a panel of self peptides measured by SPR at 37°C. **(A)** to stimulate WT c259 TCR-T cells. Target cell killing was measured after 20 hours. **(B)** The binding affinity of the c259 TCR to the peptides that induced T cell activation were measured. (Top) Representative SPR sensograms depicting injections of increasing concentrations of the c259 TCR. (Bottom) Representative equilibrium curves of c259 TCR binding to different self pMHCs. The TCR/pMHC affinity was calculated by constraining Bmax (dashed line) or fitting Bmax (solid line). **(C)** Steady-state binding affinity for the selected peptides. Barplot represents mean K_D_ ± SDs. The affinities were calculated by constraining Bmax to the value obtained from the standard curve in (B) based on the amount of BBM.1 antibody that bound the chip surface (see Methods for details). All data fitting was performed using a one site-specific binding model in GraphPad Prism.

**Figure S9:**
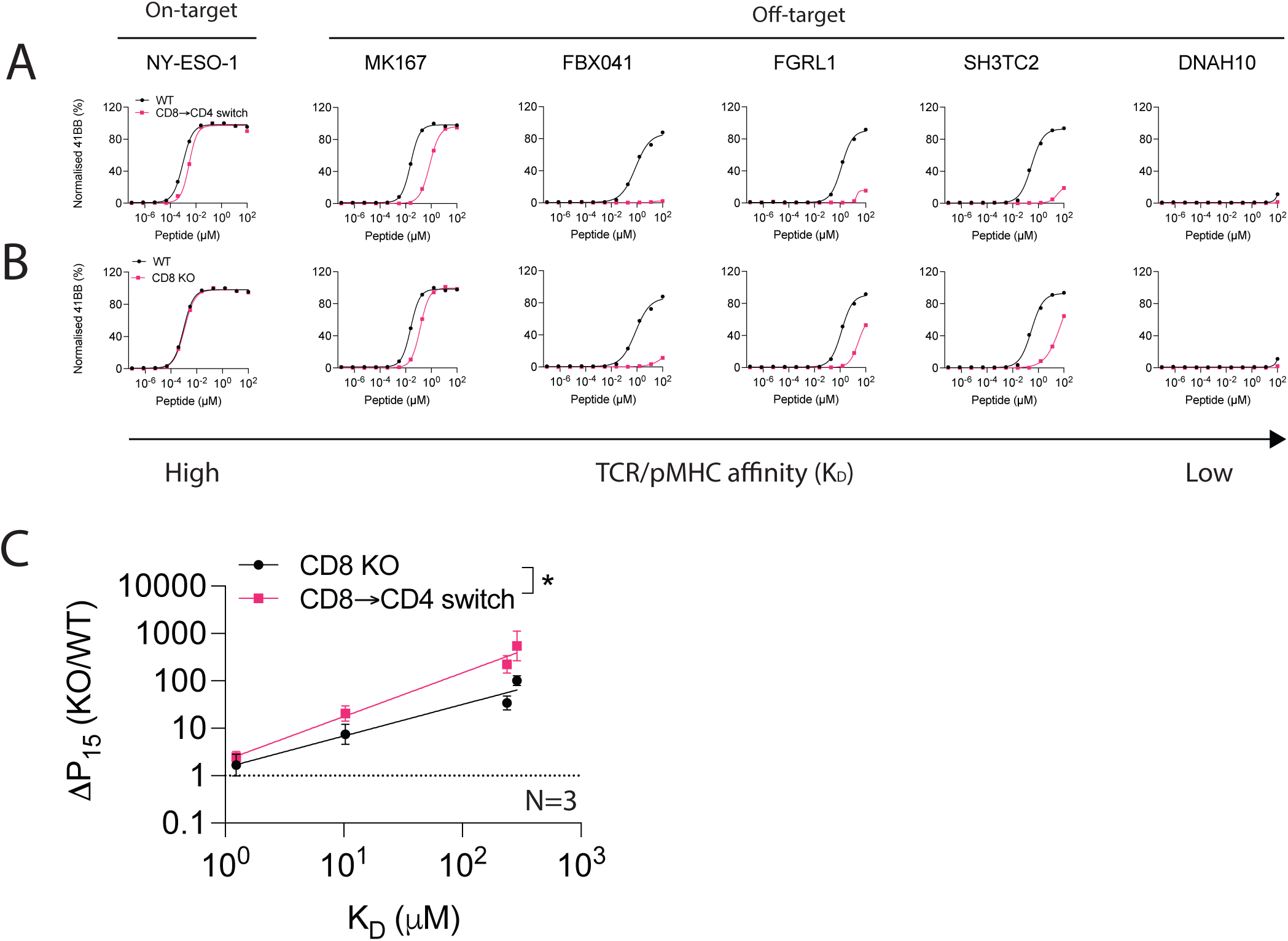
CD8→CD4 co-receptor switch cytotoxic display reduced activation against predicted self-peptides (4-1BB). **(A)** U87 cells were titrated with each of the predicted self-peptides to stimulate WT or CD8 KO cytotoxic c259 TCR-T cells. 4-1BB expression was measured after 20 hours. **(B)** U87 cells were titrated with each of the predicted self-peptides to stimulate WT or CD8→CD4 co-receptor switch cytotoxic c259 TCR-T cells. 4-1BB expression was measured after 20 hours. **(C)** Fold change in potency (P15) between modified or WT T cells from (A and B) is plotted over TCR/pMHC affinity (K_D_). Data is shown as means ± SDs. Data in (A) and (B) are representative of N=3 independent experiments with different blood donors. P values were determined by an F-test. *p*<*0.05.

## Supplementary Tables

**Table S1:** The c259 TCR affinities to the NY-ESO-1 peptide variants.

| Abbr | Sequence | Mean | SD | N |
| --- | --- | --- | --- | --- |
| 9V | SLLMWITQV | 1.240 | 0.129 | 3 |
| 6V | SLLMWVTQV | 3.538 | 0.283 | 2 |
| 3Y | SLYMWITQV | 10.350 | 1.634 | 4 |
| 6T | SLLMWTTQV | 27.628 | 4.598 | 11 |
| 4D | SLLDWITQV | 41.777 | 4.703 | 7 |
| 4A | SLLAWITQV | 62.319 | 10.313 | 8 |
| 5Y | SLLMYITQV | 157.666 | 23.818 | 3 |

**Table S2:**
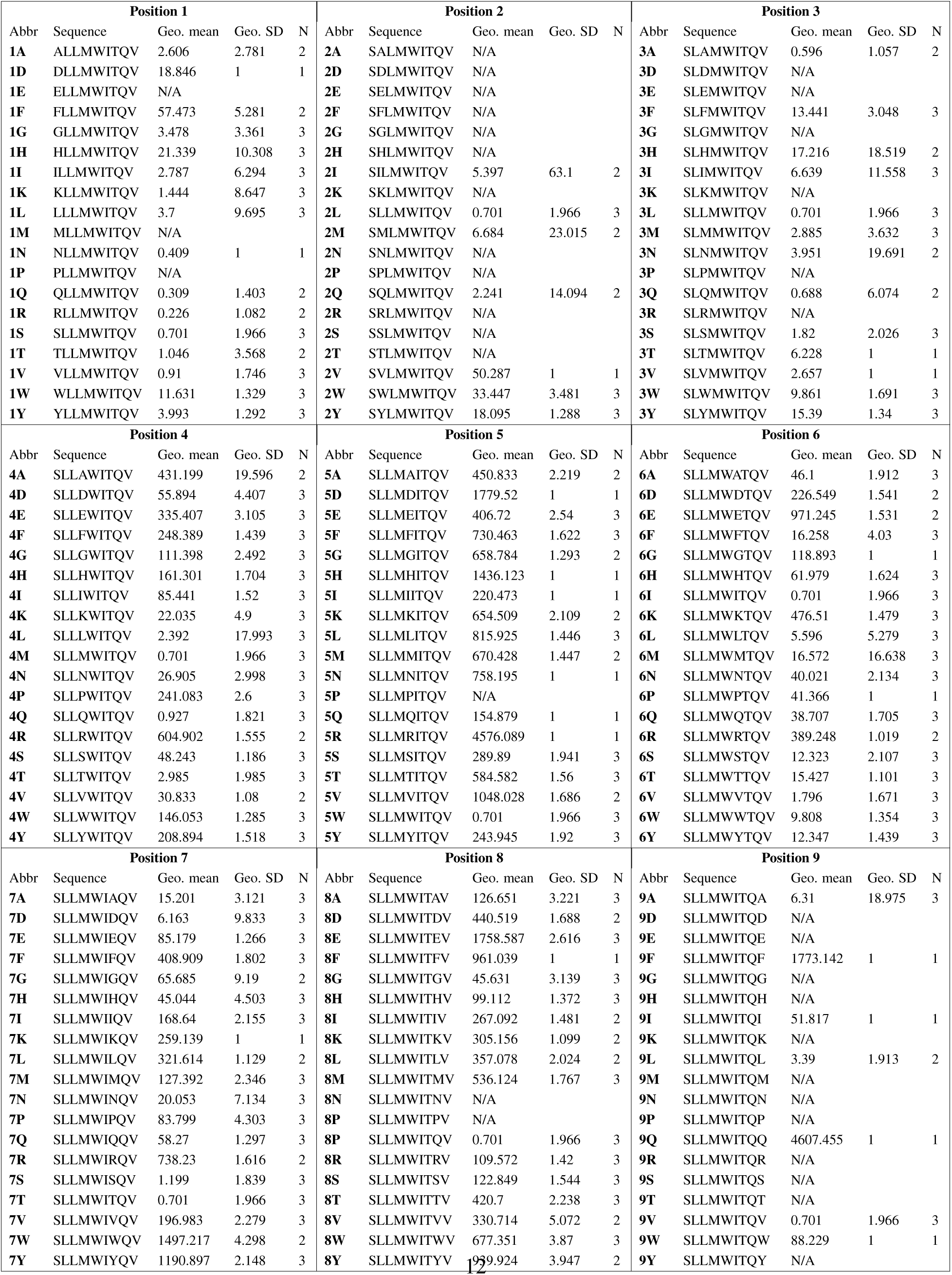
The c259 TCR affinities to the positional scanning peptide library. Geometric mean, geometric standard deviation across N experiments is reported. K_D_ values have been excluded if pMHC was unstable (indicated as N/A) or no TCR binding response was observed (indicated as NB).

**Table S3:**
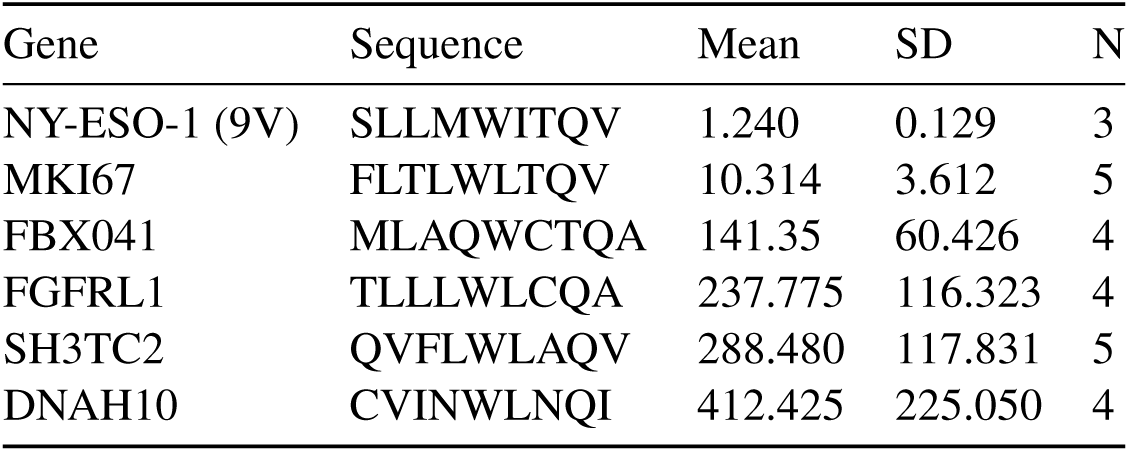
The c259 TCR affinities to predicted self-peptides.

